# Functional spreading of hyperexcitability induced by human and synthetic intracellular Aβ oligomers

**DOI:** 10.1101/2020.10.16.332445

**Authors:** Eduardo J. Fernandez-Perez, Braulio Muñoz, Denisse A. Bascuñan, Christian Peters, Nicolas O. Riffo-Lepe, Maria P. Espinoza, Peter J. Morgan, Caroline Filippi, Romain Bourboulou, Urmi Sengupta, Rakez Kayed, Jérôme Epsztein, Luis G. Aguayo

**Affiliations:** Laboratory of Neurophysiology, Department of Physiology, Universidad de Concepción, Barrio Universitario s/n P. O. Box 160-C, Concepción, Chile. Fernandez-Pérez EJ, Muñoz B, Bascuñan Muñoz DA, Peters C, Espinoza MP, Riffo Lepe NO; Institute of Neurobiology of the Mediterranean Sea (INMED), BP13 - 13273 Marseille cedex 09 - France. Morgan P, Filippi C, Bourboulou R, Epsztein J; Mitchell Center for Neurodegenerative Diseases, University of Texas Medical Branch, Galveston, TX, USA; Department of Neurology, Neuroscience and Cell Biology, University of Texas Medical Branch, Galveston, 77555 TX, USA, Sengupta U, Kayed R

**Keywords:** AMPA-R, functional spreading, hyperexcitability, Intracellular Aβ, nitric oxide

## Abstract

**Background:** Intracellular amyloid-beta oligomers (iAβo) accumulation and neuronal hyperexcitability are two crucial events at early stages of Alzheimer’s disease (AD). However, to date, no mechanism linking them has been reported.

**Methods:** Here, the effects of human AD brain-derived (h-iAβo) and synthetic (iAβo) peptides on synaptic currents and action potential (AP) firing were investigated in hippocampal neurons *in vitro, ex vivo* and *in vivo*.

**Results:** Starting from 500 pM, iAβo rapidly increased the frequency of synaptic currents and higher concentrations potentiated the AMPA receptor-mediated current. Both effects were PKC-dependent. Parallel recordings of synaptic currents and nitric oxide (NO)-related fluorescence changes indicated that the increased frequency, related to pre-synaptic release, was dependent on a NO-mediated retrograde signaling. Moreover, increased synchronization in NO production was also observed in neurons neighboring those dialyzed with iAβo, indicating that iAβo can increase network excitability at a distance. Current-clamp recordings suggested that iAβo increased neuronal excitability via AMPA-driven synaptic activity without altering membrane intrinsic properties.

**Conclusion:** These results strongly indicate that iAβo causes functional spreading of hyperexcitability through a synaptic-driven mechanism and offer an important neuropathological significance to intracellular species in the initial stages of AD, which include brain hyperexcitability and seizures.

## 1. Background

Numerous studies have reported that amyloid beta (Aβ) plays an important role in the synaptic dysfunction observed in Alzheimer’s disease (AD) patients [1,2]. Unfortunately, current therapeutic targets are focused on extracellular Aβ accumulation at a stage when the disease is well underway. However, there might be previous events in the pathology that are more important in the early pre-clinical stages, thus, other mechanisms for the pathogenesis of AD need to be considered. For example, it was recently reported that prior to the formation of extracellular Aβ deposits and intracellular tau tangles in the human brain, there is an intracellular accumulation of Aβ oligomers (iAβo) during AD, especially in vulnerable regions such as the entorhinal cortex and hippocampus [3]. Moreover, it has been suggested that amyloid plaques can result from iAβo accumulation, indicating that extracellular deposits are not the exclusive origin of senile plaques [4], thus highlighting the importance of iAβo as an early stage in the progression of the pathology. The presence of iAβo has also been associated with synaptic dysfunction in different AD mice models, which could play a key role in the cognitive deficit observed in this disease. For example, the 3xTg AD model develops intraneuronal accumulation of Aβ between 3 and 4 months of age [5], a time when cognitive deficits are first detected and a stage with little, if any, presence of extracellular Aβ [6]. Interestingly, the removal of iAβo with immunotherapy improves cognition in this model [6], and as the pathology reemerges, there is a reappearance of intraneuronal Aβ followed by the formation of extracellular amyloid plaques [7]. Thus, growing evidence supports intraneuronal accumulation of Aβ as an early event in the course of the pathology, and there are studies in at least 7 other murine AD models (including 5xFAD, APP/tau, APP751SL/PS1 KI, APPE693Δ and TBA2) [8–10] that also show the presence of intraneuronal Aβ prior to extracellular accumulation, independent of the mutation they carry.

Early intracellular Aβ accumulation is particularly interesting since recent evidence has shown that changes in neuronal excitability could also be playing a fundamental role in the initial stages of Aβ pathology, predisposing the development of a synaptopathy postulated in the initial stages of AD. For example, subjects diagnosed in the early stages of prodromal AD (a very early form of AD) displayed increased neuronal activity in the hippocampus and cortex [11,12]. Additionally, it was found that patients might exhibit increased neuronal excitability and even present a higher risk of seizures [13–15]. Indeed, two independent studies demonstrated that seizures were present in ~10-20% of patients diagnosed with sporadic AD [16,17], and patients with familial AD with mutations in APP, PS1 and PS2 also exhibited an increased prevalence of seizures [18–20]. Moreover, these early changes in excitability in human AD brains are in agreement with *in vitro* studies that showed epileptiform activity [21] and hyper-synchronous neuronal activity following acute exposure to extracellular Aβ, as well as *in vivo* models that exhibit altered intrinsic excitability [22–24] and convulsive neuronal activity [23,25]. The nature of this hyperexcitability is currently unknown.

Since intraneuronal accumulation is an early event in the pathology, these studies suggest a potential link between iAβo and neuronal hyperexcitability. However, a potential cellular mechanism that might explain this potential association has not been reported. Here, we describe that h-iAβo and iAβo applied to the post-synaptic neuron increased global synaptic transmission, excitability, and neuronal synchronization in different *in vitro* and *in vivo* hippocampal models. The presynaptic action of iAβo was primarily caused by a NO-dependent retrograde signaling, while the postsynaptic effect was mediated by the potentiation in AMPA currents by a PKC-dependent mechanism.

## 2. Methods

Detailed description of the applied methods are provided in Additional file 1: Supplementary methods

### 2.1. Primary cultures of rat hippocampal neurons

Hippocampal neurons were obtained from 18-day embryos from pregnant Sprague-Dawley rats and cultured for 10-14 days *in vitro* (DIV) as previously described [94].

### 2.2. Preparation of amyloid beta oligomers

Human Aβ42 fluorescently labeled with FAM (5 (6) – Carboxyfluorescein) at the N-terminal or without fluorescence were bought from Biomatic (USA) and Genemed Synthesis Inc. (USA), respectively. Synthetic oligomeric species of Aβ (later referred as iAβo) were prepared as previously described [95]. See details in Additional file 1: Supplementary methods, section 1.1.

### 2.3. Obtaining human-derived Aβo

Oligomeric assemblies of Aβ were derived from AD brain tissues (later referred as h-iAβo) by following previously published protocol [96]. Aβo was extracted from the PBS-soluble fraction of AD brain homogenate using co-Immunoprecipitation Kit (ThermoFisher Scientific, USA) following manufacturer’s guidelines. Briefly, amine-reactive resin was coupled with anti-Aβ 6E10 antibody (BioLegend, USA) followed by incubation with PBS-soluble fraction of AD brain homogenate. Bound proteins were eluted in 0.1 M glycine (pH 2.8), and the final pH was adjusted to 7.0 by adding 1 M Tris-HCl (pH 8). Eluted fraction was subjected to buffer exchange and collected in sterile PBS. This fraction was further separated by size exclusion chromatography using AKTA Explorer system fitted with a Superdex 200 Increase 10/300 GL Column. Degassed PBS was used as mobile phase with a flowrate of 0.5 mL/min to collect the Aβo fraction. The total protein concentration was measured with bicinchoninic acid protein assay (Pierce™ Micro BCA Kit, ThermoFisher Scientific, USA). Human brain-derived Aβo was characterized by Western blot analysis and atomic force microscopy (details in Additional file 1: Supplementary methods, section 1.2 and 1.3).

### 2.4. Electrophysiology

Whole cell recordings were used to simultaneously apply Aβ peptide intracellularly through the internal solution contained in the recording electrode and record postsynaptic currents at constant voltage (voltage-clamp) or action potentials (current-clamp) *in vitro, ex vivo* and/or *in vivo*.

#### 2.4.1. Voltage-clamp experiments *in vitro* and *ex vivo*

For *in vitro* experiments, the dish culture medium was replaced with a normal external solution (NES) containing (in mM): 150 NaCl, 5.4 KCl, 2.0 CaCl2, 1.0 MgCl2, 10 glucose and 10 HEPES (pH 7.4 adjusted with NaOH, 310 mOsm / L). Cells were stabilized at room temperature for 20 minutes before beginning the experiments. Unless otherwise noted, the internal solution used to record voltage-clamp experiments, including synaptic currents or ligand-evoked currents, contained (in mM): 120 KCl, 2.0 MgCl2, 2 Na2ATP, 10 BAPTA, 0.5 NaGTP and 10 HEPES (pH 7.4 adjusted with KOH, 290 mOsm / L). To study miniature post-synaptic currents (mPSCs), 500 nM tetrodotoxin (TTX) (Alomone Labs, Israel) was applied in the NES of the well containing the cells (details in Additional file 1: Supplementary methods, section 1.5). For the AMPAergic evoked (eEPSCs) and GABAergic (eIPSCs) currents, 500 nM of TTX was used in the recording solution, and using a gravity-driven perfusion system (2-3 ml/min), 100 μM of extracellular AMPA and GABA were applied, respectively. All currents (synaptic and evoked) were recorded in voltage-clamp mode by adjusting the membrane potential to -60 mV (unless otherwise noted). Voltage-clamp *in vitro* experiments were performed using an Axopatch-200B amplifier (Molecular Devices, USA) and an inverted microscope (Nikon Eclipse TE200-U, Japan). The acquisition was made using a computer connected to the registration system using a Digidata 1440A acquisition card (Molecular Devices, USA) and the pClamp10 software (Molecular Devices, USA). Electrodes with a resistance of 4-5 MΩ were pulled from borosilicate capillaries (WPI, USA) in a horizontal puller (P1000, Sutter Instruments, USA). Some voltage-clamp experiments were performed using the following internal solution (in mM): 114 K-Gluconate, 4 KCl, 4 MgCl2, 10 BAPTA and 10 HEPES (pH 7.4 adjusted with KOH, 290 mOsm / L). This was later referred as “*low Cl^-^ internal solution*”. Details of voltage-clamp *ex vivo* experiments in Additional file 1: Supplementary methods, section 1.5.

#### 2.4.2. Current-clamp experiments *in vitro*

To study the membrane potential (Vm) recordings were made in current-clamp mode as previously described [97], using the previously mentioned *low Cl^-^ internal solution*. To evoke action potentials, a family of current pulses applied for 300 ms was used (from -300 pA to +275 pA, increasing by 25 pA steps). In some experiments, the voltage-dependent sodium channel intracellular inhibitor (QX-314) (Tocris, USA) was used. Before starting the recording of evoked action potentials, a small holding current (−2 to - 50 pA) was applied to stabilize the resting membrane potential (RMP) to -70 mV. Current-clamp *in vitro* experiments were performed using an Axopatch-200B amplifier (Molecular Devices, USA) and an inverted microscope (Nikon Eclipse TE200-U, Japan).

#### 2.4.3. Current-clamp recordings *in vivo*

All experiments on live animals were approved by the Institut National de la Santé et de la Recherche Médicale (INSERM) Animal Care and Use Committee, in accordance with the guidelines of the European Community Council directives (2010/63/EU). Data were obtained from male Wistar rats between the ages of postnatal day 25 (P25) to P35 (weight range, 90-110 g). Recordings were made as previously described [98]. See details in in Additional file 1: Supplementary methods, section 1.6.

### 2.5. Histology

At the end of the recording period and to confirm the location and the morphology of *in vivo* recorded neurons, we performed immunohistochemical analysis of hippocampus sections. See details in Additional file 1: Supplementary methods, section 1.7.

### 2.6. Simultaneous recordings of electrophysiology and fluorescence

Simultaneous studies were performed using the same methodology described for the recording of synaptic currents in voltage-clamp mode *in vitro* (section 2.4.1) together with NO fluorescence using a previously described methodology [99]. See details in Additional file 1: Supplementary methods, section 1.8.

### 2.7. Data analysis

All data obtained from all parameters were plotted using OriginPro2019b (Origin Lab, USA). Data are shown as mean ± SEM for normally distributed populations and as median and interquartile ranges (IQR) for non-normally distributed populations. Statistical analyses were performed using the two-tailed unpaired Student’s t-tests (α=0.05) or the two-tailed Mann-Whitney U-test (α=0.05) as appropriate, after testing for normality with Shapiro-Wilk test (for n<50) or Kolmogorov-Smirnov (for n>50) and for homogeneity of variances with Levene’s test. Data with more than two groups or factors were analyzed by two-way ANOVA test (α=0.05). One-way ANOVA test was used to compare several populations of neurons, followed by Tukey or Welch’s ANOVA with Games-Howell post-hoc test to correct for variance heterogeneity using R software [100] (www.r-project.com). A probability level (p) < 0.05 was considered statistically significant (* p < 0.05, ** p < 0.01, *** p < 0.001). Unless otherwise noted, all “n” values represent recorded cells (see details of parameter calculations in Additional file 1: Supplementary methods, section 1.9).

## 3. Results

### Rapid effect of human derived iAβo in hippocampal neuronal activity

We examined the synaptic effects of human AD-brain derived intracellular oligomers (h-iAβo) using an electrophysiological approach. First, characterization of the biochemically purified Aβ species with WB analysis using A11 (anti-amyloid oligomers) and 6E10 (generic anti-Aβ) antibodies confirmed the presence of Aβo in this preparation (Fig. 1a) and AFM imaging showed a homogeneously distributed population of sphere shaped oligomers ranging from 5-20 nm in size (Fig. 1b). Using a patch electrode, we delivered increasing concentrations (0, 50 and 1000 nM) of h-iAβo while recording post-synaptic currents in primary hippocampal neurons (see scheme in Fig. 1c). Control synaptic current recordings were made with an intracellular solution without h-iAβo. Electrophysiological recordings showed the presence of spontaneous synaptic events (Fig. 1d, arrows in blue) that had a stable response over a period of 10 minutes. Applying a low concentration of h-iAβo with the patch electrode (50 nM) augmented the presence of bursts of spontaneous synaptic currents interspersed in the recording (Fig. 1d, arrowheads in middle trace). It is also possible to observe that part of the total activity is mediated by spikes in current recording mode, which also augmented with the presence of 50 nM h-iAβo (Fig. 1d, arrows in middle trace). The increase in activity was a concentration-dependent phenomenon, given that there were more bursts of synaptic activity and events of greater amplitude as the concentration of iAβo augmented from 50 to 1000 nM. We integrated the area under the current trace for each condition and computed the charge transferred for the recorded cell (see calculation details in Additional file 1: Supplementary methods, section 1.9). 1000 nM h-iAβo increased significantly the charge transferred meaning that more current by unit of time was flowing through the membrane compared to control conditions (Fig. 1e). Interestingly, 50 and 1000 nM produced an increase in the frequency of the recorded events (Fig. 1f), but only 1000 nM augmented the amplitude (Fig. 1g). To confirm the time course of these results and to validate the method as an approximation to effectively deliver Aβo to the intraneuronal compartment, we repeated this experiment using a fluorescently labeled synthetic Aβo. The results show that Aβo in the patch pipette is rapidly delivered to the intraneuronal compartment, since the increase in synaptic activity was correlated with a temporal increase in fluorescence, with a t_1/2_ of 29 s (Additional file 1: Supplementary Figure 1).

**Figure 1.**
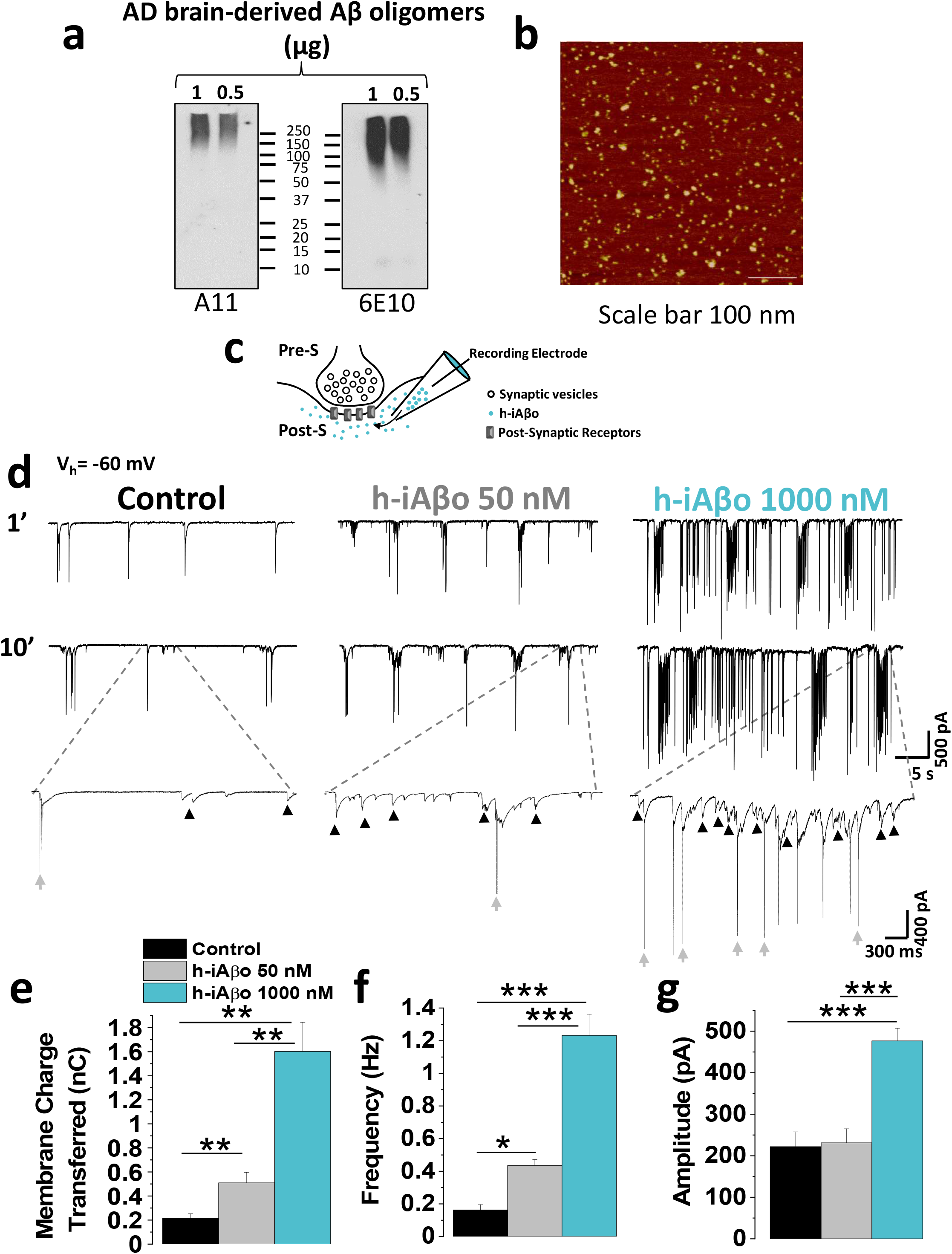
Intracellular human derived Aβ oligomers (h-iAβo) increased the synaptic transmission and AP spike firing in hippocampal neurons *in vitro*. **a,** Western blot analysis of AD brain-derived Aβ oligomers with A11 and generic Aβ antibody 6E10. **b,** AFM image of AD brain-derived Aβ oligomer. Scale bar 100 nm. **c,** Schematic representation of the synaptic recordings, showing the pre-synaptic (Pre-S) and post-synaptic (Post-S) compartment, and the use of the patch electrode to dialyze the cell with h-iAβo and to record membrane currents (electrode do not represent actual size). **d,** Representative synaptic currents recordings (holding potential (Vh) = -60 mV). Bursts of synaptic currents (arrowheads) and spikes in current recording mode (arrows) are observed. **e-g,** Quantification of charge transferred (**e**) frequency (**f**) and amplitude (**g**) of post-synaptic currents. Bars represent the average ± SEM for control (n=6), h-iAβo 50 nM (n=7) and h-iAβo 1000 nM (n=8) cells. One-Way Welch’s ANOVA with Games-Howell post-hoc test for (**e**): F(2,9.91)=24.04, p=1.57E-4. p-values for post-hoc test: control vs. h-iAβo 50 nM: 5.68E-3, control vs. h-iAβo 1000 nM: 1.51E-9 and h-iAβo 50 nM vs. h-iAβo 1000 nM: 6.76E-3. One-Way Welch’s ANOVA with Games-Howell post-hoc test for (**f**): F(2,10.69)=32.82, p=2.73E-5. p-values for post-hoc test: control vs. h-iAβo 50 nM: 3.29E-2, control vs. h-iAβo 1000 nM: 1.17E-4, h-iAβo 50 nM vs. h-iAβo 1000 nM: 3.80E-4. One-Way Welch’s ANOVA with Games-Howell post-hoc test for (**g**): F(2,11.09)=19.25, p=2.47E-4. p-values for post-hoc test: control vs. h-iAβo 1000 nM: 6.94E-04, h-iAβo 50 nM vs. h-iAβo 1000 nM: 4.67E-4. * denotes p <0.05, ** p <0.005, *** p <0.001.

### Concentration-dependent increase of miniature currents by iAβo

It is accepted that in order to record synaptic currents at the postsynaptic site, a release of neurotransmitters must occur from the pre-synaptic terminal (Fig. 1c). To quantify the effect of iAβo on actual synaptic transmission, we recorded miniature post-synaptic currents (mPSCs). Intracellular application of 500 nM of h-iAβo was able to produce a rapid and significant increase in the frequency and amplitude of mPSCs (Fig. 2a - c). This result was confirmed using different concentrations of synthetic iAβo, that compared with control condition, exhibited a significant increase in the number of events as the concentrations was raised, indicating a concentration-dependent effect (Fig. 2d). The effects of iAβo were more evident on the frequency of miniature events since statistically significant differences were already found at 0.5 nM of iAβo when compared to control conditions (Fig. 2e). On the other hand, the effect of iAβo on the amplitude was only significant at a much higher concentration (Fig. 2f, 1000 nM). We wanted to perform some additional experimental controls to evaluate potential confounds with the application of intracellular iAβo. Therefore, we applied reverse sequence Aβo, as well as the vehicle used to dissolve the peptide. Under these conditions, no significant differences in frequency with respect to the control condition were found (Additional file 1: Supplementary Figure 2). Furthermore, we wanted to control whether the osmolarity of the internal solution was affected by adding different iAβo concentrations, but no significant changes were found (Additional file 1: Supplementary Figure 2). Finally, we applied iAβo in the absence and presence of a specific antibody (A11) that recognizes oligomeric forms [26] and found that it significantly attenuated its toxicity (Additional file 1: Supplementary Figure 3).

**Figure 2.**
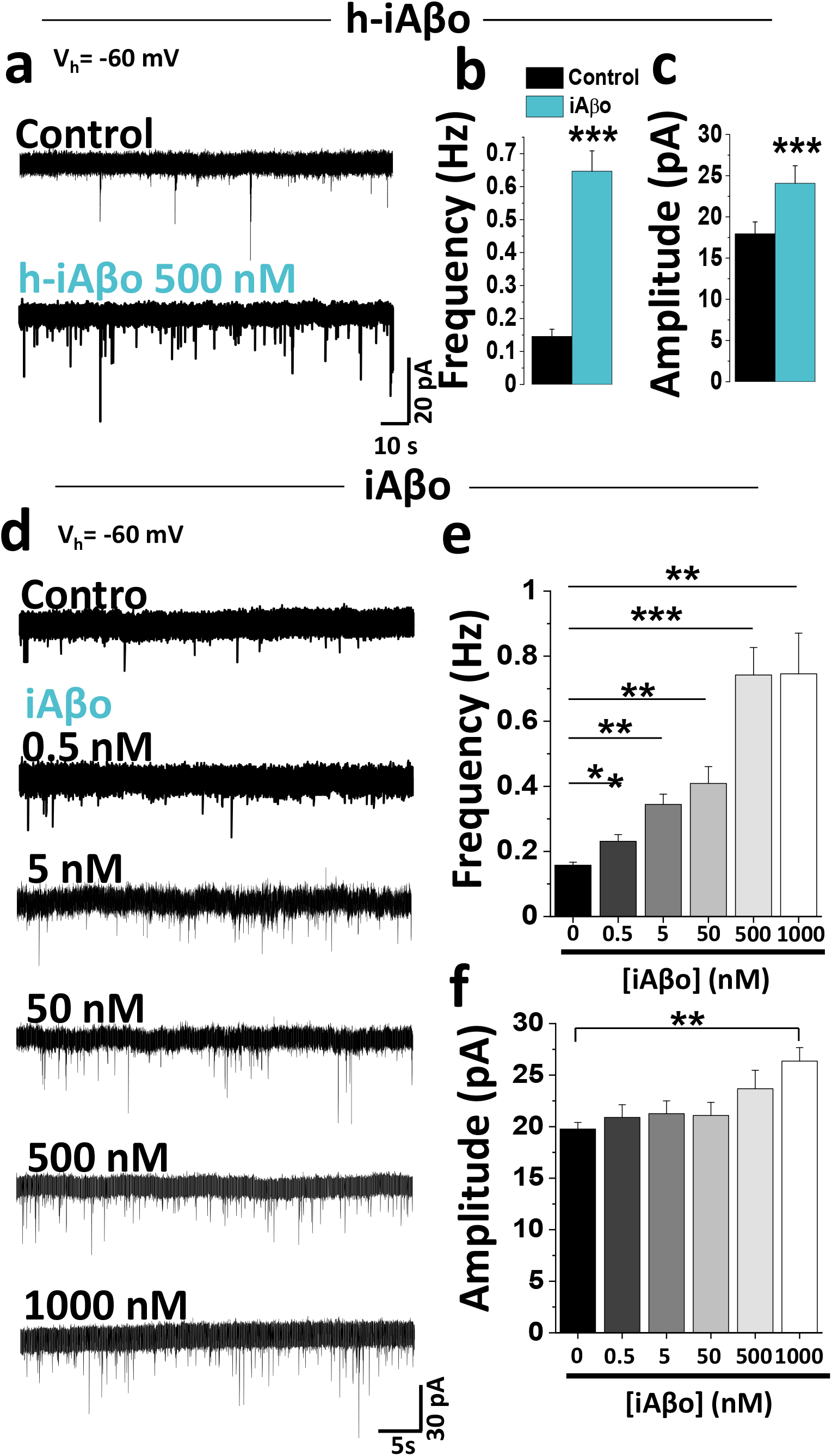
Effects of intracellular Aβ oligomers on the frequency and amplitude of miniature post-synaptic currents. **a,** Representative mPSCs traces after applying h-iAβo 0.5 μM (Vh = -60 mV). **b, c,** mPSCs frequency (**b**) and amplitude (**c**) quantification for control (n=5 cells) and h-iAβo 500 nM (n=9 cells). **d,** mPSCs traces for control (n=16) and after applying increasing concentrations of synthetic iAβo 0.5 nM (n=14), 5 nM (n=16), 50 nM (n=16), 500 nM (n=14) and 1000 nM (n=16 cells). **e, f,** mPSCs frequency (**e**) and amplitude (**f**) for each of the conditions described in (**d**). Bar charts represent the average ± SEM. Unpaired Student’s t-test for (**b**) (t(12)=-5.845, p=7.89E-05) and unpaired Student’s t-test with Welch’s correction for (**c**) (t(11.93)=-5.123, p=2.56E-04). One-Way Welch’s ANOVA with Games-Howell post-hoc test for (**e**): F(5,86)=13.76, p= 7.15E-10. p-values for post-hoc test: iAβo 0 nM vs. 0.5 nM: 4.57E-2, iAβo 0 nM vs. 5 nM: 2.73E-4, iAβo 0 nM vs. 50 nM: 2.33E-3, iAβo 0 nM vs. 500 nM: 1.17E-4, iAβo 0 nM vs 1000 nM: 3.11E-3. One-Way Welch’s ANOVA with Games-Howell post-hoc test for (**f**): F(5,86)=3.73, p=0.004. p-values for post-hoc test: iAβo 0 nM vs. 1000 nM: 2.01E-3. * denotes p <0.05, ** p <0.005 and *** p <0.001.

### Human-iAβo affects excitatory/inhibitory (E/I) balance *in vitro*

A great body of evidence indicates that Aβ disrupts excitatory neurotransmission in different AD mice models [27–30] thereby affecting E/I balance [31]. Since our previous results showed an increase in iAβo-mediated neurotransmission and augmented spike number in current recording mode, we examined whether h-iAβo could specifically affect glutamatergic vs GABAergic neurotransmission and thereby changing the E/I balance in hippocampal neurons. For this, we performed voltage-clamp experiments using a low Cl^-^ internal solution (see details in section 2.4.1). This allowed us to record EPSC and IPSC separately by changing the membrane holding potential. The results showed that h-iAβo markedly increased the spontaneous excitatory post-synaptic currents (sEPSC) (Fig. 3a, bottom, right trace), but not the inhibitory post-synaptic currents (sIPSC). The data also shows that charge transferred for sEPSC, but not sIPSC, was significantly increased by h-iAβo (Fig. 3b, black bar). Consequently, the E/I balance obtained analyzing charge transferred was highly and rapidly increased by h-iAβo (Fig. 3b, white bar). Since sEPSC are mainly mediated by AMPA receptors in the current experimental conditions, our results suggest that the alteration in AMPA-mediated neurotransmission underlies this excitatory input disruption.

**Figure 3.**
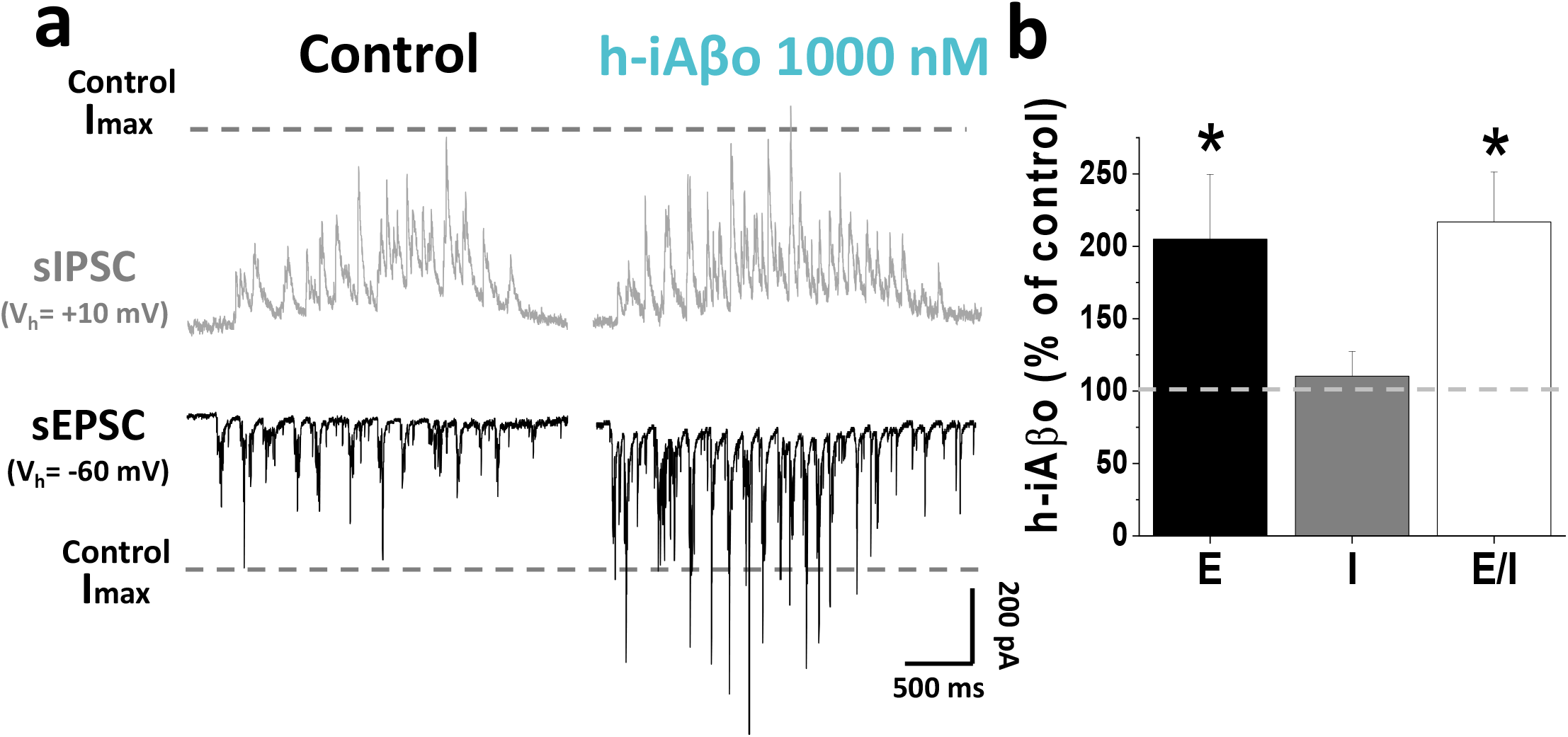
h-iAβo induces a shift in excitatory/inhibitory (E/I) balance. **a,** Spontaneous inhibitory (sIPSC) and excitatory (sEPSC) post-synaptic recordings obtained in absence (control) and presence of 1 μM h-iAβo. In this conditions, reverse potential for Cl^-^ was ≈ -66 mV, allowing us to record sEPSC at a V_h_ of -60 mV and sIPSC at +10 mV. Segmented lines show the maximum amplitude observed for sEPSC and sIPSC in control condition (Control I_MAX_). **b,** Charge transferred for sEPSC (E) (unpaired Student’s t-test: t(11)=-2.75, p=1.86E-2), charge transferred for sIPSC (I) and E/I ratio for each condition (unpaired Student’s t-test: t(11)=-2.73, p=1.93E-2), expressed as percentage of control condition. Bar charts represent the average ± SEM for control (n=7) and h-iAβo (n=6) cells. * denotes p <0.05.

### Differential effects of iAβo on isolated excitatory AMPA and inhibitory GABA_A_ currents

Subsequently, we studied pharmacologically isolated AMPAergic and GABAergic miniature synaptic currents (2 of the predominant neurotransmissions present in hippocampal neurons [32]) to confirm the selectivity of iAβo. In the presence of iAβo, there was a significant increase in the frequency of AMPA mEPSCs and GABAergic mIPSCs (Fig. 4a and c). On the other hand, only the amplitude of AMPA was affected by iAβo, as clearly seen by the average synaptic event (Fig. 4b). The quantification of these parameters (frequency and amplitude) were plotted to assess the iAβo effects on both types of neurotransmission (Fig. 4e and f). We also performed voltage-clamp recordings in the stratum pyramidale of the CA1 area (SP-CA1) in hippocampal slices of an adult mouse (*ex vivo*) and found similar results (Additional file 1: Supplementary Figure 4). Because the AMPAergic mediated-neurotransmission was more significantly affected, we examined AMPAergic currents in presence of h-iAβo and confirmed that human-derived oligomers also produced the increase in excitatory synaptic transmission (Additional file 1: Supplementary Figure 5). The experimental observation that the amplitude was affected for AMPA receptors (with human-derived and synthetic preparations) was also confirmed by evaluating ligand-evoked currents in hippocampal neurons. The results showed that the presence of iAβo enhanced the AMPA-mediated current amplitude by about 2 times (Control I normalized: 1.00 ± 0.05 vs. iAβo: 1.95 ± 0.23) (Fig. 4g and i). Once again, this was a specific effect for the excitatory transmission since no effects were observed for GABA-evoked currents (Fig. 4h and i).

**Figure 4.**
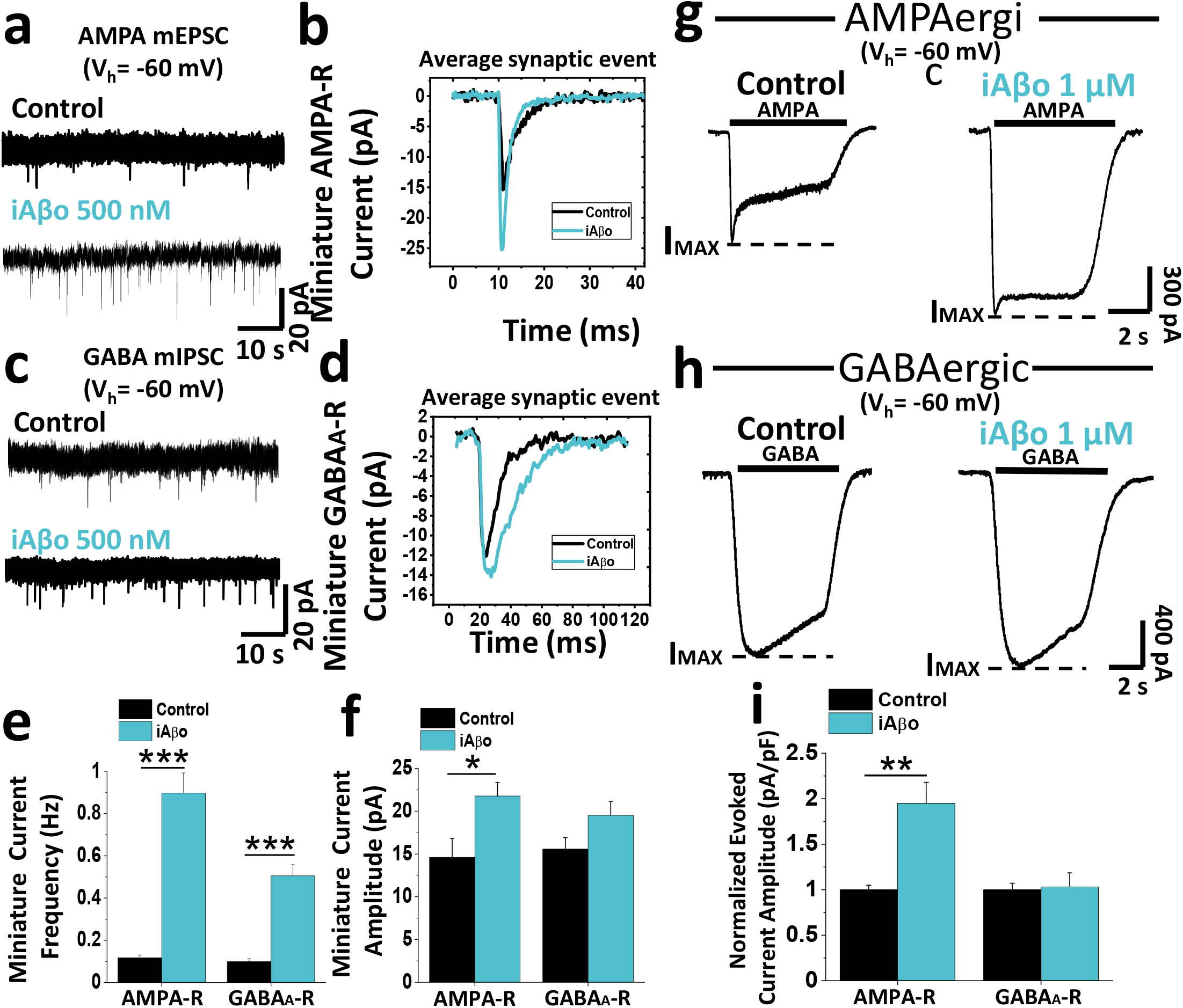
iAβo differentially affected AMPA-R and GABA_A_-R mediated miniature and evoked PSCs. **a - d,** Pharmacologically isolated AMPA (**a**) and GABA (**c**) mPSCs in control condition and with iAβo 500 nM at Vh = -60 mV, showing the averaged synaptic current for each condition (**b** and **d**). **e, f,** mEPSC (n=9) and mIPSC (n=8) frequency (**e**) and amplitude quantification (**f**) for control and iAβo-treated cells. **g, h,** AMPA (**g**) and GABA (**h**) evoked currents in the absence (n=9) and presence of iAβo 1 μM (n=9). Segmented lines indicate maximum current reached (I_MAX_). **i,** Normalized evoked current amplitude (value corresponding to I_MAX_) for AMPA and GABA receptors. Bar charts represent the average ± SEM. Unpaired Student’s t-test with Welch’s correction for mEPSC frequency: t(8.29)=-8.07, p=3.31E-5 and unpaired Student’s t-test for mIPSC frequency: t(14)=-7.43, p=3.20E-6. Unpaired Student’s t-test for mEPSC amplitude: t(16)=-2.62, p=0.018 and mIPSC amplitude: t(14)=-1.86, p=0.086. Unpaired Student’s t-test with Welch’s correction for (**i**) AMPA-R: t(16)=-3.83, p=1.4E-3 and GABA_A_-R: t(16)=0.17, p=0.863. * denotes p < 0.05, ** p <0.005 and *** p <0.001.

### The effect of iAβo on AMPA neurotransmission depends on PKC

These previous results prompted us to investigate in depth the mechanism(s) that could be mediating the effect of iAβo on the AMPAergic excitatory transmission. It has been shown that phosphorylation of ionotropic channels plays a preponderant role in the regulation of synaptic function [33], and the intracellular perfusion of an atypical isoform of PKC denominated PKCM (a constitutively active form of PKC) increases the synaptic response mediated by the AMPA receptor in a similar way to what we previously observed in our recordings with iAβo [34]. Taking into account the above, and considering that it is quite a rapid effect, we decided to record AMPA-R currents (miniature and ligand-evoked) in the presence of a PKC inhibitor (CLR, 2.5 μM intracellularly). We found that the effect of iAβo on the frequency and amplitude of the mEPSCs was diminished when co-applied together with CLR (Fig. 5a), as evidenced by quantifying both parameters for all experimental conditions tested with and without inhibitor application (Fig. 5b and c), clearly showing that the effect of iAβo on AMPA mEPSCs was dependent on this kinase. On the other hand, AMPA evoked-currents also showed that iAβo increased the current evoked by AMPA (Fig. 5d) in relation to control, but in the presence of the PKC-inhibitor a significant decrease occurred, reaching values similar to those of control where only CLR was applied (Fig. 5d). Quantification of the amplitude of the evoked EPSC showed a statistically significant increase in presence of iAβo with respect to control conditions(Fig. 5e). This was interesting because it not only demonstrated that PKC was involved in the effect of iAβo on the amplitude of mEPSCs and evoked EPSC (a post-synaptic effect), but also on the frequency of mEPSCs (pre-synaptic effect), suggesting that the global effects of iAβo on the excitatory synapse through this kinase compromises the pre- and post-synaptic compartments.

**Figure 5.**
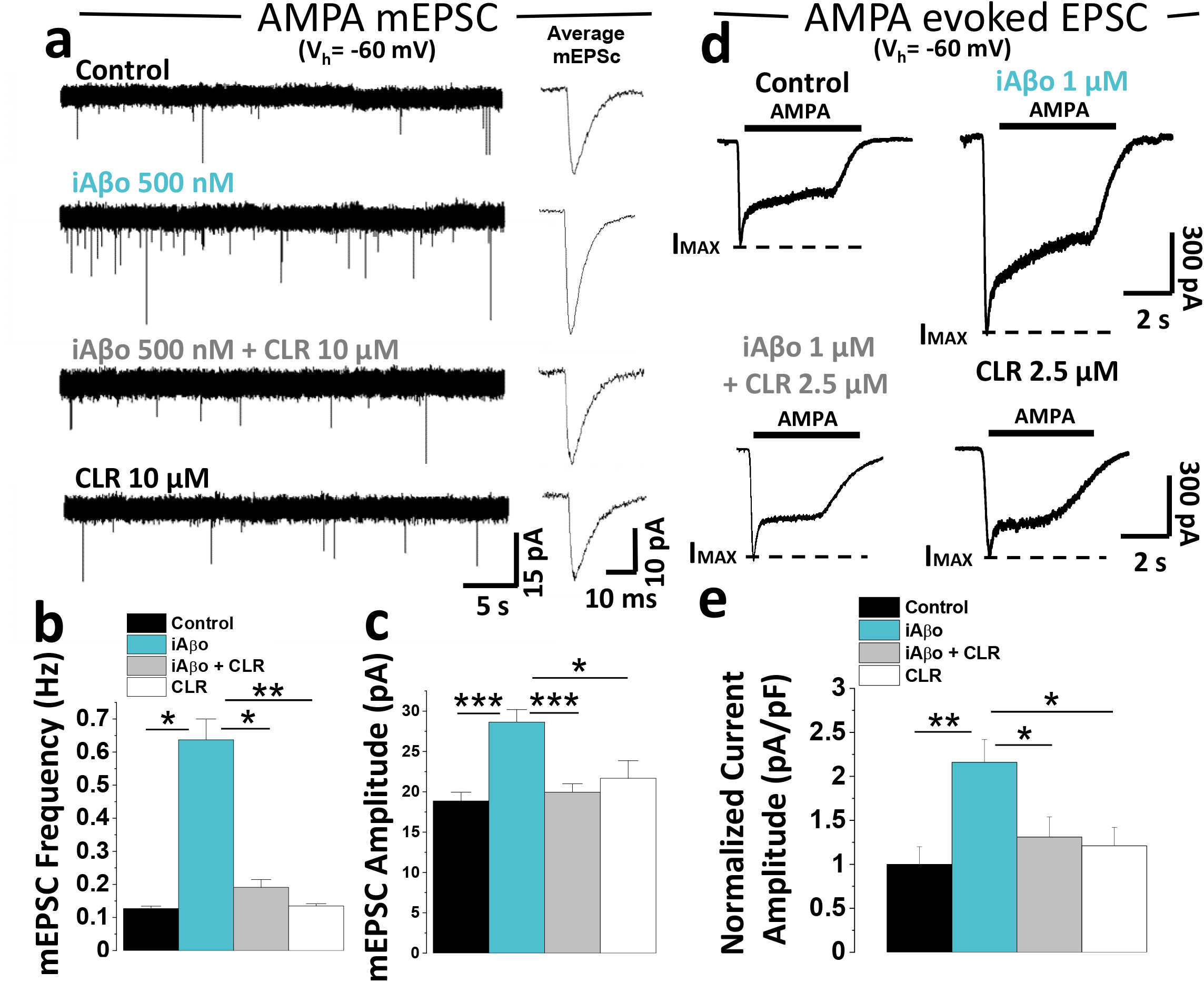
The effect of iAβo on AMPAergic miniature and evoked EPSCs is PKC dependent. **a,** mEPSCs recordings obtained for control (n=11), iAβo 500 nM (n=11), chelerythrine (CLR) 2.5 μM + iAβo 500 nM (n=12) and CLR 2.5 μM (n=12) (Vh = -60 mV). **b, c,** mEPSC frequency (**b**) and amplitude quantification (**c**) for each of the conditions described in (**a**). **d,** AMPA evoked current in the absence (n=9) and presence of iAβo 1000 nM (n=9), iAβo co-applied together with CLR (n=11) and CLR (n=11) (Vh = -60 mV). Segmented lines indicate maximum current reached (I_MAX_). **e,** Average amplitude of AMPA evoked current for each condition in (**d**). Bar charts represent the average ± SEM. One-Way Welch’s ANOVA with Games-Howell post-hoc test for (**b**): F(3,42)=6.11, p=0.002. p-values for post-hoc test: control vs iAβo: 2.27E-2, iAβo vs. iAβo + CLR: 1.78E-2, iAβo vs. CLR: 2.07E-3. One-Way Welch’s ANOVA with Games-Howell post-hoc test for (**c**): F(3,42)=8.54, p=1.52E-4. p-values for post-hoc test: control vs. iAβo: 2.65E-4, iAβo vs. CLR: 1.23E-2, iAβo vs. iAβo + CLR: 9.49E-4. One-Way Welch’s ANOVA with Games-Howell post-hoc test for (**e**): F(3,35)=5.88, p=2.31E-3. p-values for post-hoc test: control vs. iAβo: 3.31E-3, iAβo vs. CLR: 1.10E-2, iAβo vs. iAβo + CLR: 1.49E-2. * denotes p < 0.05, ** p <0.005 and *** p <0.001.

### Increased NO-production in iAβo-treated in recorded (RN) and the neighboring (NN) neurons

The data showed that there was a potent effect on the frequency of mPSCs when iAβo was applied to the postsynaptic neuron, suggesting an increase in the release of neurotransmitters at the pre-synaptic terminal [35]. Could it be that iAβo applied to the postsynaptic neuron affects the release of neurotransmitter from the pre-synaptic neuron? If so, could there be a retrograde mechanism involved in the effect? [36]. Because the effect was rapid, we thought that it could be produced by nitric oxide (NO), a retrograde messenger, known to be involved in presynaptic neurotransmitter release [37–39] and produced by nitric oxide synthase (NOS). It is important to consider that low concentrations of iAβo were used for all the recordings obtained in these experiments, which did not affect mPSCs amplitude, but only its frequency. Under these conditions, we decided to use a non-selective nitric oxide synthase inhibitor (L-NAME) together with the following methodological approach: mPSCs and relative NO levels using a fluorescent probe DAQ were simultaneously recorded (details in >section 2.6 and in Additional file 1: Supplementary methods, section 1.8). In addition, not only the fluctuation of the relative levels of NO in the recorded cell (denoted as RN) (Fig. 6a) was quantified, but also of neighboring neurons (denoted as NN) (Fig. 6a). Pre-incubation of hippocampal neurons with L-NAME showed no significant differences in the frequency of miniature currents (Fig. 6b-d), but did significantly decrease the NO levels in RN (Fig. 6e) and in NN (Fig. 6g) when compared to control conditions. The statistical comparison was performed at the end of the recording period (20 min.) (Fig. 6e and g, indicated by the gray box) and plotted as a bar graph for RN (Fig. 6f) and NN (Fig 6h). For its part, iAβo had the expected effect and increased the frequency of mPSCs (Fig. 6b-d). Interestingly, the increase in iAβo-mediated miniature synaptic activity occurred at the same time that NO levels increased in the RN (Fig. 6e-f). More importantly, iAβo not only increased the levels of NO in the RN, but also in NN that did not get iAβo (Fig. 6g-h). Pre-incubation of the neuronal culture with L-NAME reduced the effect of iAβo on mPSCs frequency by ≈ 51% (Fig. 6b-d). This decrease in the frequency of mPSCs correlated well with the decrease in the relative NO levels in the RN and NN (Fig. 6e-h, iAβo + L-NAME). Additionally, this NO-dependent effect was confirmed by co-applying iAβo together with a donor molecule of NO (SNAP 300 μM) causing a synergistic effect between the two, increasing the frequency of mPSCs in the RN (Additional file 1: Supplementary Figure 6), as well as in the relative levels of NO in both the RN and NN (Additional file 1: Supplementary Figure 6). The opposite effect was observed using iAβo in the presence of the NO chelator C-PTIO (Additional file 1: Supplementary Figure 6). On the other hand, no changes in the effects of iAβo in the frequency of mPSCs were found using a specific inhibitor of inducible NO (iNOS) (Additional file 1: Supplementary Figure 7).

**Figure 6.**
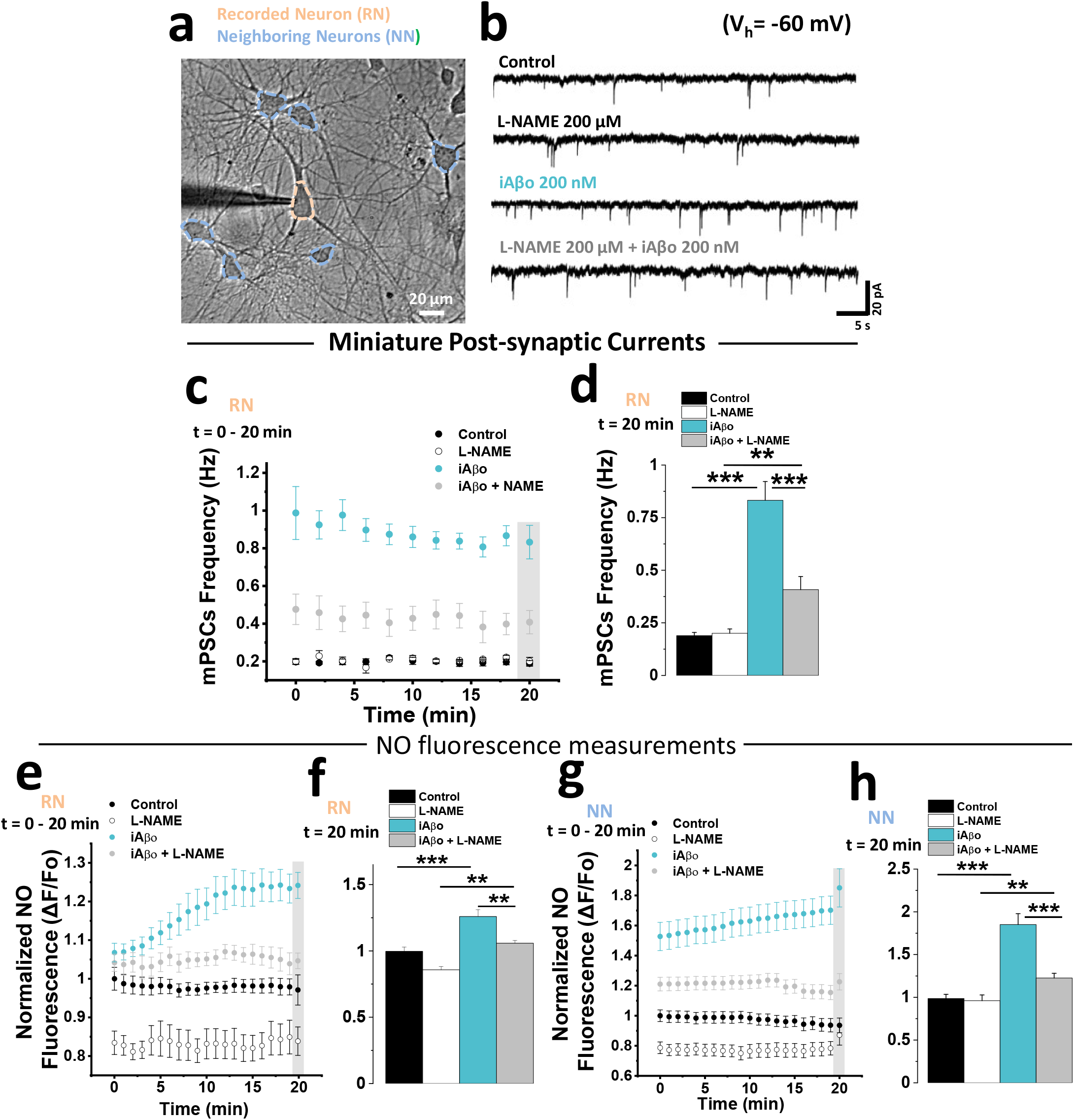
Nitric oxide synthase is important the effects of iAβo on the frequency of miniature synaptic currents in the neuronal ensemble. **a,** Micrograph showing the area of the culture in which mPSCs were obtained from the recorded neuron (RN), as well as the NO-fluorescence for RN and of the neighboring neurons to RN (NN). **b, c,** Representative recordings (**b**) and quantification of mPSCs frequency (**c**) obtained during the course of 20 minutes in absence (control) and presence of iAβo 200 nM, together with the pre-incubation with L-NAME (200 μM for 20-30 min) (Vh = -60 mV). **d,** Quantification of mPSCs frequency at 20 min (obtained from data marked inside gray rectangle from plot in **c**). **e - h,** Relative levels of NO (expressed as normalized fluorescence) obtained throughout the course of the experiment and at 20’ for RN (**e** and **f**, respectively) and NN (**g** and **h**, respectively). Line and bar graphs represent the average ± SEM. Control (n=9), L-NAME (n=8), iAβo (n=11) and iAβo + L-NAME (n=10) for RN and control (n=97), L-NAME (n=102), iAβo (n=119) and iAβo + L-NAME (n=102) for NN. One-Way Welch’s ANOVA with Games-Howell post-hoc test for (**d**): F(3,34)=60.309, p=1.07E-13. p-values for post hoc test: Control vs. iAβo: 1.06E-08, L-NAME vs. iAβo + L-NAME: 7.80E-3, iAβo vs. iAβo + L-NAME: 1.40E-08. One-Way Welch’s ANOVA with Games-Howell post-hoc test for (**f**): F(3,34)=21.926, p=4.41E-8. p-values for post hoc test: Control vs. iAβo: 2.53E-05, L-NAME vs. iAβo + L-NAME: 3.03E-3, iAβo vs. iAβo + L-NAME: 1.18E-03. One-Way Welch’s ANOVA with Games-Howell post-hoc test for (**h**): F(3,416)=40.763, p=4.11E-23. p-values for post hoc test: Control vs. iAβo: 6.44E-20, L-NAME vs. iAβo + L-NAME: 2.16E-3, iAβo vs. iAβo + L-NAME: 8.85E-09. ** denotes p <0.005, *** p <0.001.

### iAβo increased the excitability of hippocampal neurons *in vivo* and *in vitro*

The results showed that the excitatory transmission and E/I balance are strongly affected by human and synthetic iAβo. The CA1 area in the hippocampus (a major center of excitatory neurotransmission) is one of the first areas prone to develop early AD neuropathology in humans and mice models [40–44], therefore, we decided to record hippocampal CA1 pyramidal neurons using whole cell current-clamp recordings in an anesthetized *in vivo* rat model to examine neuronal excitability (Fig. 7a). We found that in the presence of iAβo, the neuron begun spiking in response to lower current stimuli (Fig. 7b). Plotting the number of spikes vs. injected current showed a strong shift of the curve to the left (Fig. c), implying that the cell with iAβo was more excitable. Indeed, the constant rheobase for firing was diminished in neurons treated with iAβo (Fig. 7d). On the other hand, the calculated values for input resistance (R_in_) and kinetic parameters of action potentials (APs) (amplitude, duration and threshold) in both conditions were similar to control conditions (Additional file 1: Supplementary Figure 8). Furthermore, the recording of resting membrane potentials (Vm), without current injection, evidenced a large increase in Vm fluctuations when iAβo was applied to the neuron (Fig. 7e). Vm fluctuations generated an assortment of local potentials that had more depolarized values in iAβo-treated neurons (Control: -68.02 ± 0.001 v/s iAβo: -63.55 ± 0.004; Fig. 7f) nearing it to the AP firing threshold, which was also evidenced as a significant change in Vm standard deviation (SD) (Fig. 7g). iAβo also increased the chance of spontaneous spike firing (Fig. 7h) (control: 0.003 ± 0.003 Hz vs iAβo: 1.068 ± 0.463 Hz). Post recording immunohistochemical analysis confirmed the location of the recorded neurons in the stratum pyramidale of dorsal hippocampus at CA1 (Fig. 7i – j).

**Figure 7.**
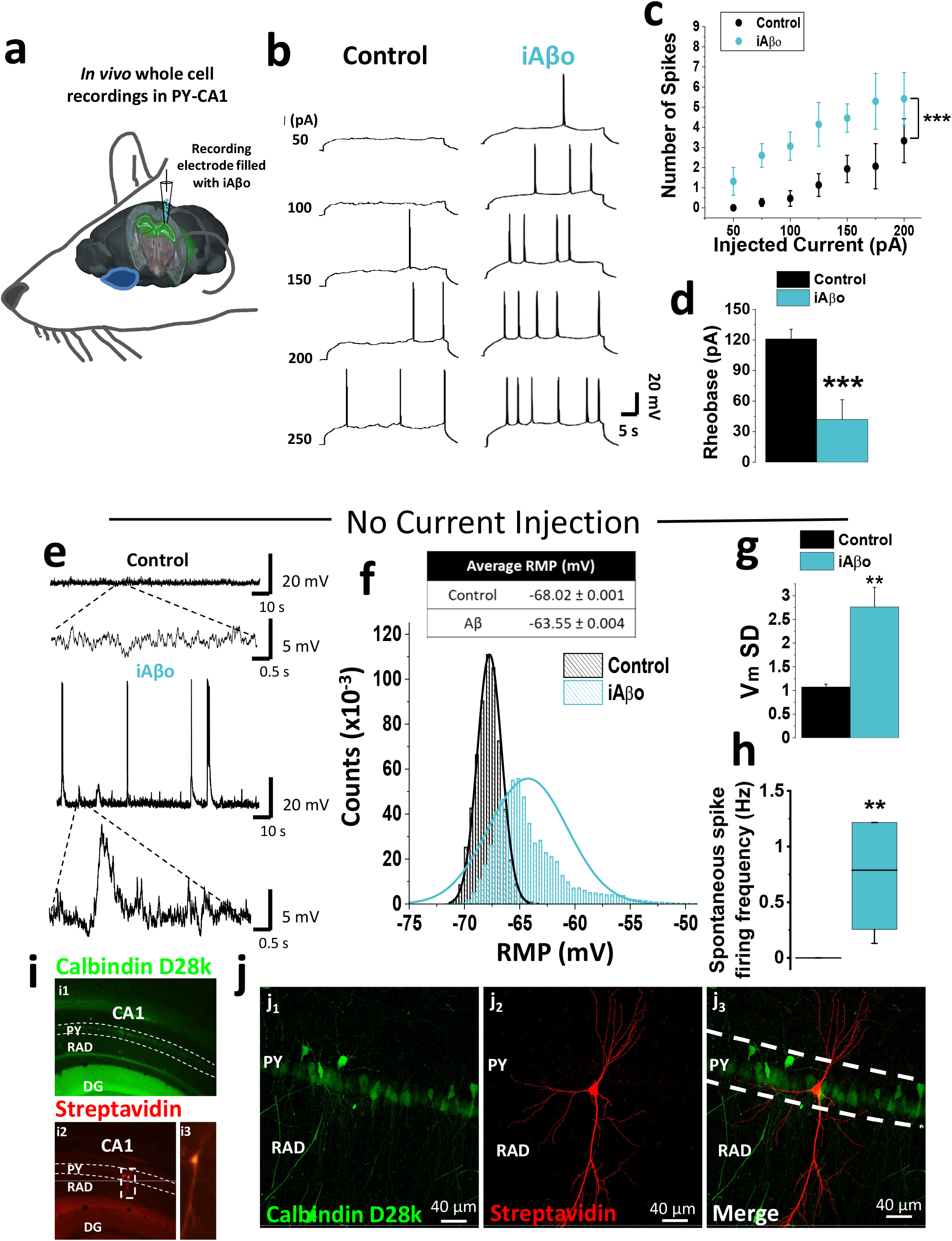
iAβo increased the firing of action potentials evoked by current injection in hippocampal neurons *in vivo*. **a,** Scheme representing the location of the electrode filled with iAβo to record action potentials in the pyramidal cell layer of CA1 (PY) of rat hippocampus (highlighted in green) (brain image obtained from Allen Institute Brain Atlas). **b,** Representative action potential recordings in absence and presence of 500 μM iAβo. **c,** Relationship between the number of evoked action potentials and the injected current intensity for the experimental conditions described in **b** (two-way ANOVA: F(7,103)=7.983, p=1.01E-7). **d,** Quantification of the rheobase constant (unpaired Student’s t-test: t(14)=4.188, p=9.11E-4). **e,** Representative recordings obtained without injection of depolarizing current pulses after stabilizing the resting membrane potential (RMP) to -70 mV. **f,** Vm values histogram and average values of Vm ± SEM. **g, h,** Quantification of standard deviation (SD) values for Vm (**g**) (unpaired Student’s t-test: t(14)=-3.445, p=3.94E-3) and spontaneous AP firing frequency (**h**) (Mann-Whitney U-test: U=0, z-score=-3.199, p=1.38E-3). **i,** Epifluorescence micrographs of coronal cuts obtained after the electrophysiological recording, showing a positive staining for calbindina-D28k (in green, **i_1_**) and streptavidin (in red, **i_2_**), in hippocampus-CA1. The recorded neuron (**i_3_**) is also observed. DG, dentate gyrus; PY, pyramidal cell layer of CA1; RAD, stratum radiatum. **j,** Confocal micrographs, demonstrating the positive staining for streptavidin in the recorded neuron (in red, **j_2_**) as well as the merge with calbindina-D28k (**j_3_**) within the stratum pyramidale of the CA1 region of the hippocampus. Bar and line charts represent the average ± SEM for control (n=10) and h-iAβo (n=6) cells of at least 6 rats. ** denotes p <0.005, *** p <0.001.

After confirming the effects on excitability in an *in vivo* model and with the aim of examining the disease relevance and to contribute mechanistically to the results, we examined neuronal excitability hippocampal neurons. For this, current clamp experiments were performed and the effects of human derived (h-iAβo) and synthetic (iAβo) oligomers were evaluated. In agreement with the previous data, we found that a smaller current injection was required to trigger the firing of action potentials (AP) in the presence of h-iAβo (Fig. 8a). Plotting the number of spikes vs. injected current demonstrated that the stimulus-response curve shifted to the left indicating an increase in neuronal excitability in the presence of h-iAβo (Fig. 8b) together with a reduced rheobase (Fig. 8c). Similar results were obtained for synthetic iAβo in the same model (Additional file 1: Supplementary Figure 9), confirming the effect of human and synthetic preparations on neuronal excitability *in vitro*.

**Figure 8.**
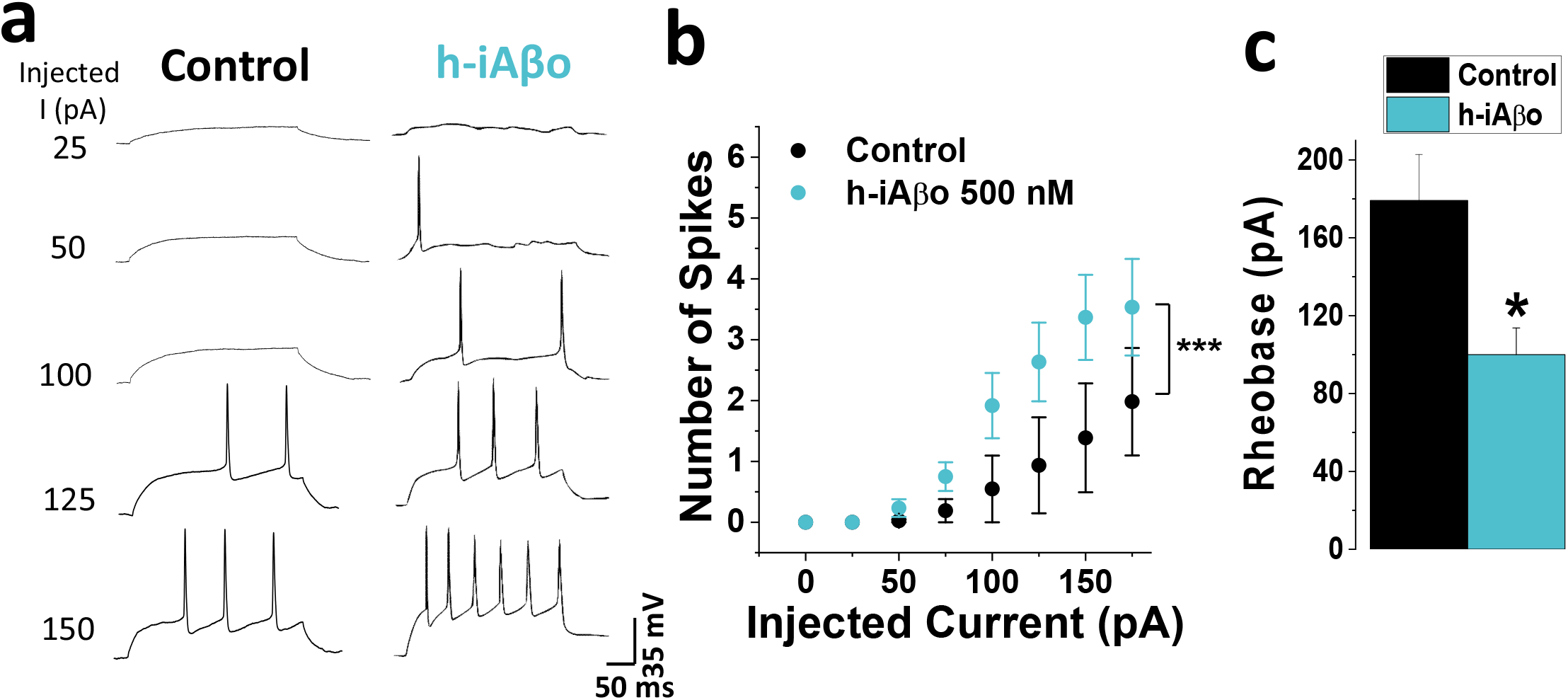
Intracellular human derived Aβ oligomers (h-iAβo) increased the firing of action potentials in hippocampal neurons *in vitro*. **a,** Hippocampal neuron action potential (AP) recordings in the absence and presence of 500 nM h-iAβo. **b,** Relationship between the number of AP spikes and the injected current intensity for the experimental conditions described previously (two-way ANOVA: F(8,79)=9.077, p=9.31E-9). **c,** Rheobase constant decreased ≈45% for h-iAβo condition (unpaired Student’s t-test: t(9)=2.737, p=2.29E-2). Bars and line charts represent the average ± SEM for control (n=6) and h-iAβo (n=5) cells. * denotes p <0.05, *** p <0.001.

We then evaluated if the effect of h-iAβo on action potentials (APs) could be the result of augmented transitory depolarizations at the post-synaptic level due to the increase in AMPAergic neurotransmission. In order to examine this, we applied h-iAβo and recorded the membrane potential (Vm) fluctuations of hippocampal neurons without current injection in the presence and absence of an intracellular voltage-dependent sodium channel (Na_v_) blocker (5 mM QX-314). When Nav were blocked, APs were not evident in the recorded neuron. In control conditions, Vm fluctuations generated a population of local synaptic potentials (Fig. 9a) with an average Vm value of -65.3 ± 0.003 mV (Fig. 9b), while in the presence of h-iAβo there was a significant increase in the number and amplitude of these depolarizing events (Fig. 9a), with an average of -60.6 ± 0.012 mV (Fig. 9b), similar to what we observed previously in the *in vivo* model. In the presence of iAβo and QX-314 no major differences were observed in Vm fluctuations, except for the lack of AP spikes (Fig. 9a), showing a similar average Vm value to the one observed when h-iAβo was present (−60.8 ± 0.017 mV) (Fig. 9b). On the other hand, Vm in the presence of QX-314 had a similar average value to control condition with an average of -65.1 ± 0.014 mV (Fig. 9a-b, in gray). Similar results were obtained in the presence of synthetic iAβo (Additional file 1: Supplementary Figure 10). It is interesting to note that in all tested conditions the external application of an AMPA receptor antagonist (CNQX) through the perfusion system attenuated all Vm fluctuations (Fig. 9a), indicating that these changes in membrane potential in our experimental condition were synaptic AMPA-driven potentials. This confirms that the AMPAergic-mediated synaptic input is responsible for the iAβo-mediated hyperexcitability observed in our experiments.

**Figure 9.**
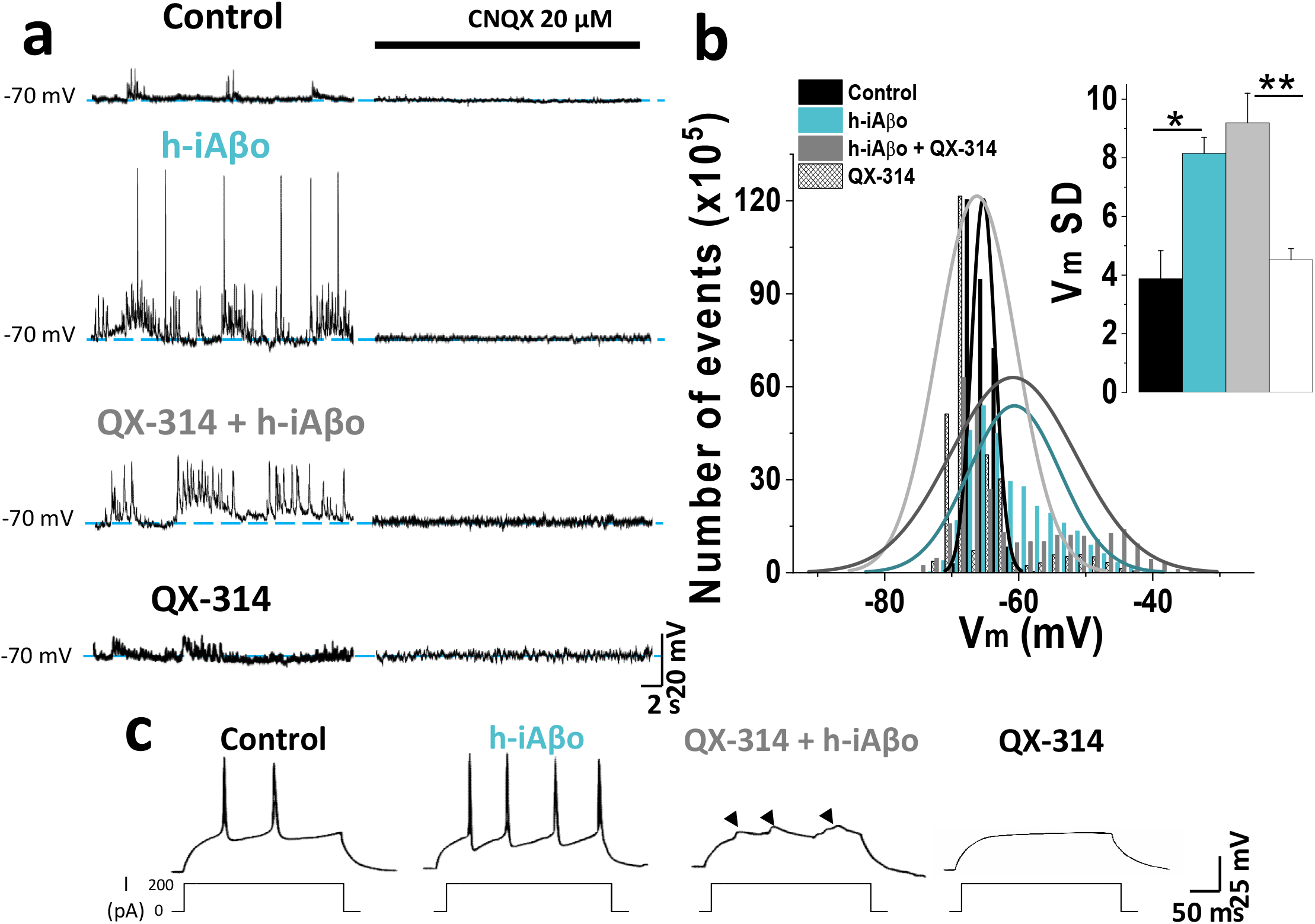
Intracellular blockade of voltage-regulated Nav channels does not prevent depolarization of the membrane activated by h-iAβo. **a,** Representative recordings obtained without current injection, showing membrane potential (Vm) fluctuations under the different conditions tested. External application of CNQX 20 μM inhibited the fluctuations at great extent in all conditions. **b,** Histogram showing the distribution of Vm values in the different experimental conditions shown in **a**, along with the Vm SD (inset bar graph) One-Way Welch’s ANOVA with Games-Howell post-hoc test for: F(3,19)=11.299, p=1.77E-4. p-values for post hoc test: Control vs. iAβo: 7.13E-3 and QX-314 vs. QX-314 + iAβo: 2.12E-3). **c,** Current injection experiments demonstrating that, under the control and h-iAβo conditions, the generation of action potentials was not inhibited, while Na_v_ intracellular block by QX-314 prevented spiking of neurons with h-iAβo and without it. Arrowheads over 3th trace (left-to-right direction) indicate the presence of depolarizing post-synaptic potentials when h-iAβo was present. This did not occurred for the condition with QX-314 alone (4th trace; left-to-right direction). Bars represent the average ± SEM for control (n=6), h-iAβo (n=5), h-iAβo+QX-314 (n=6) and QX-314 (n=6) cells. * denotes p < 0.05, ** p <0.005 and *** p <0.001.

In presence of intracellular QX-314, no spikes (APs) were detected (Fig. 9c, 3^th^ and 4^th^ traces; left-to-right direction) confirming blockade of Na+ channels. We also observed the presence of transitory depolarization (Fig. 9c, arrowheads over 3^th^ trace; from left to right) that were not present in presence of QX-314 alone (Fig. 9c, 4^th^ trace; from left to right). This data indicates that iAβo exerts depolarizations of the post-synaptic membrane even in the absence of APs, which are mediated by AMPA receptors, suggesting that this effect did not depend on the generation of APs at the post-synaptic level, but rather an increase in synaptic transmission at pre- and post-synaptic levels.

In order to rule out the possibility that iAβo was directly affecting intrinsic membrane properties and AP firing, we used a model of dorsal root ganglion neurons (DRG) that do not form synapses *in vitro* to record action potentials in the presence and absence of iAβo. The results showed that iAβo did not affect the AP firing nor did the rheobase constant (Additional file 1: Supplementary Figure 11a-d), contrary to the results found in hippocampal neurons. Additionally, and similar to hippocampal neurons, no significant differences were found in the parameters calculated for the APs evoked in this experiment such as amplitude, duration, and threshold (Additional file 1: Supplementary Figure 11e-g).

## 4. Discussion

Presence of oligomeric species of Aβ in the intraneuronal domain of the membrane was able to increase neurotransmission through mechanisms that involved pre-and post-synaptic actions that resulted in an increased excitability and spreading of neuronal activation. Because the present findings were obtained with human AD-brain derived and synthetic Aβo, they should contribute with disease-relevant data in a new and significant working model. Furthermore, using *in* vitro models allowed us to mechanistically understand the effects on neurotransmission and excitability that could be important in the initial stages of AD and that is likely occurring in an AD-human brain.

Contrary to previously published data using extracellular Aβ or with more chronic application models [45–50], we did not find any synaptic deficits, instead, iAβo increased excitatory and inhibitory neurotransmissions, but with more marked effects on the former. This raises the question of how could iAβo actions progress to synaptic dysfunction in the first place? In this regard, it has been reported that extracellular Aβ increases glutamate neurotransmission [51] which could lead to excitotoxicity [52–54] together with vesicular depletion over extended periods of time [55,56] [57–59]. If we consider these observations with the effects reported in this study, it is tempting to speculate that, over time, iAβo influences synaptic depletion and this is in agreement with results from murine AD models that show that early intracellular accumulation of Aβo leads to synaptic failure and the onset of cognitive deficits (for further details see [60]), once again emphasizing the importance of iAβo during the pre-symptomatic stage of AD.

### Pre-synaptic mechanism: Increase in retrograde synaptic signaling by NO

Simultaneous fluorescence and electrophysiology recordings demonstrated that NO is involved in the pre-synaptic effect of iAβo. The data showed that iAβo applied to the postsynaptic site augmented NO levels globally in the neurons together with the frequency of mPSCs, reflecting an enhanced release of neurotransmitters into the synaptic space [35], which is in agreement with studies that have shown that NO increases neurotransmitter release from the excitatory and inhibitory synapsis [61–63]. Furthermore, PKC is likely involved in this pre-synaptic effect of iAβo since CLR prevented the iAβo-mediated increase in the mEPSCs frequency (Fig. 10a). In agreement with this notion, it was reported that PKC also participates in pre-synaptic mechanisms mediated by the gas neuromodulator NO [64–66], emphasizing the pathway that iAβo might be using to increase neurotransmission.

**Figure 10.**
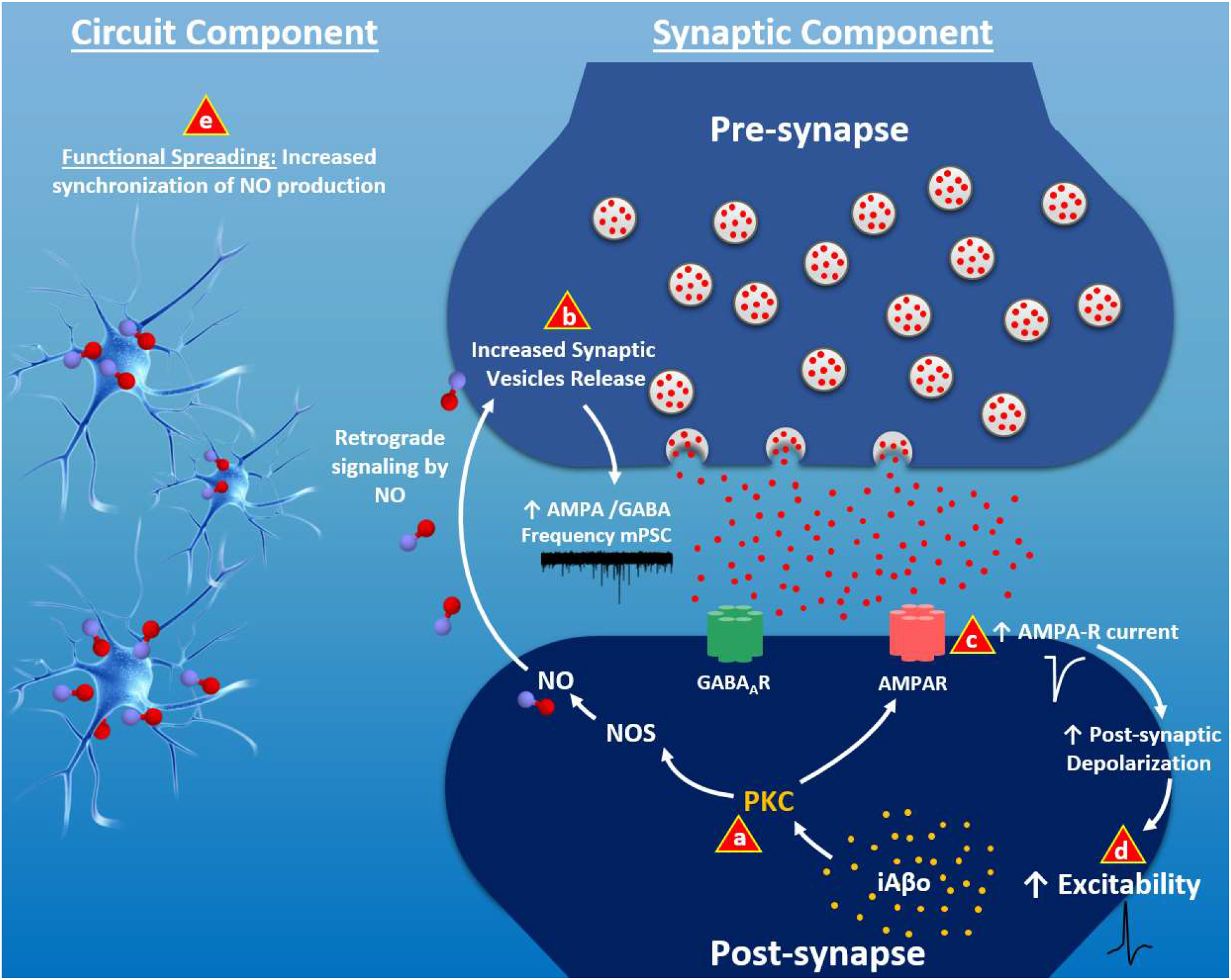
Scheme depicting the effects of intracellular effects of Aβo on neurotransmission and neuronal excitability. Schematic summary of iAβo actions at the pre- and post-synaptic levels, highlighting a synaptic and a circuit component. At synaptic level, being present in the post-synaptic compartment activates PKC (**a**), which in turn mediates the activation of nitric oxide synthase (NOS). This triggers the production of nitric oxide (NO) at the post-synaptic level, which, through a retrograde mechanism, activates an increase in the release of neurotransmitters from the pre-synaptic compartment (**b**). These activate the post-synaptic receptors AMPAR (pink) and GABA (in green), which was globally evidenced as an increase in the frequency of GABAergic and AMPAergic mPSCs, respectively. Additionally, PKC present in the post-synapse increases the AMPA receptor current (**c**), increasing post-synaptic depolarization, facilitating the approach of Vm to the threshold and generating action potentials more easily. This directly affects the neuronal function, increasing the excitability of the cell (**d**). Finally, at the circuit level, iAβo in the recorded neuron increased synchronization NO production in nearby neurons that don’t have iAβo, a phenomenon we named as “functional spreading” (**e**).

The support for NO-signaling involvement in iAβo-mediated pathology is multiple. First, we found that iAβo increased this messenger in an ensemble of neurons. Second, during the development of AD, NO synthases (NOS) increase in rodent models and in human brains [67,68]. Moreover, in pre-symptomatic 3xTg mice, it has been observed that this alterations that favor NO-synthesis occur together with synaptic pathology [68]. Thus, we can hypothesize that unregulated release of NO in AD pathology might have negative consequences for neuronal function. Third, at the cellular level a prolonged release of NO could also cause metabolic, oxidative stress, and dysfunction in organelles such as mitochondria and ER, eventually leading to neuronal death [69–73]. Therefore, the results agree with current studies indicating that NO signaling is altered through different mechanisms during the development of AD.

### Post-synaptic mechanism for AMPA transmission potentiation

The data indicate that iAβo caused a post-synaptic potentiation through the increase in the AMPA-mediated current. Generally speaking, a change in mEPSC amplitude is a reflection of a change at the post-synaptic level [74]. However, this is not the only case, since this might also be explained by an increased neurotransmitter content in synaptic vesicles, translating into an increase in the amplitude of post-synaptic currents [75]. This was not the case for iAβo because evoked currents with a saturating concentration of AMPA still increased the amplitude of the current, supporting the idea that the effect was postsynaptic in nature. Physiologically relevant, the amplitude of AMPA-evoked and miniature-AMPAergic currents (mEPSCs) increased in the presence of iAβo, an effect that was largely attenuated in the presence of a PKC inhibitor (CLR). In general, several kinases found in hippocampal neurons, including PKA (Protein kinase A) and CAMKII (Ca^2+^/Calmodulin Protein kinase II), can have similar effects may affect AMPA receptors. For example, CAMKII increases AMPA receptor conductance [76] and the number of synaptic physical contacts, improving the connectivity that mediates excitatory transmission [77]. On the other hand, PKA can increase the number of AMPAR at the synapse [78], as well as increase the release of neurotransmitters at the pre-synaptic site [79].

In this regard, Whitcomb and co-workers reported a PKA dependent increase on AMPA current amplitude in response to iAβo in hippocampal slices [80]. However, their results differed from the present because they required longer times of exposure and it was dependent on calcium. The discrepancies may be due to differences in oligomers preparation, time exposure course, or the effective concentration, since in the present study the amplitude increased only with higher concentrations. Interestingly, another study in cultured autaptic hippocampal neurons found that intracellular Aβ decreased neurotransmission by an unknown mechanism [81]. The present results allows us to conclude that the increase in AMPA receptor function represents a main mechanism by which iAβo depolarizes the post-synaptic membrane and increases the probability for action potential firing (Fig. 10c and d).

### Implications of pre- and post-synaptic effects at circuit level in AD pathology: NO functional spreading and neural excitability in AD

The question that became evident after finding that iAβo was able to produce large increases in the E/I balance was if this could have a more global consequence on neuronal hyperexcitability. First, we were able to demonstrate that iAβo induced rapid and large modifications of the bioelectric activity of hippocampal neurons leading to an “epileptiform like phenomena”, which was evidenced by the presence of complex discharge activity composed of synaptic depolarization and spike firing, both initiated by an increase in AMPA neurotransmission.

A quite novel result found during our study was that not only the hippocampal neuron that had iAβo became overactive, but also neighboring neurons displayed increases in NO-production in a synchronized fashion. This suggests that there is a phenomenon in which the functional effect of iAβo extends to surrounding neurons in a coordinated way. In other words, by mechanisms only now beginning to be appreciated, iAβo alters the function of hippocampal neurons beyond where it is physically present, thus, iAβo has an impact at the circuit level (Fig. 10e).

This phenomenon of increased synchronization, closely resembles the effect of prions of the nervous system, which have been shown to spread the pathology through the brain [82,83]. The difference is that in our case there would be no transmission of iAβo *per se*, but only of the functional effects that it produces. Therefore, we have adopted the term “functional spreading” to refer to this phenomenon.

Now, the consequence of increasing NO is functional because as NO is released the probability of increasing the discharge of excitatory neurotransmitters in nearby areas is also increased [84–86], thus increasing neuronal activity, frequency of action potential firing, and excitability of the neural network [71], [87,88]. Also, in an epilepsy model induced by the intrahippocampal injection of kainate, a well-known AMPA receptor agonist, an increase in NOS activity was observed suggesting a direct link between NO and AMPA-mediated epileptiform activity [89]. There is a strong link between AMPAR and epilepsy [90], and the antagonists significantly reduced or nullified epileptiform activity in hippocampal neurons [91,92]. Thus, our results support the existence of a strong relationship between NO and glutamatergic neurotransmission in the genesis of epileptic hyperexcitability, which could be contributing to the neuronal excitability observed in our models and to the epileptiform activity exhibited by murine AD models and more importantly AD patients [14,15,25,93].

Considering our results, it is possible to hypothesize that only a few neurons that with iAβo could initiate the increase in neurotransmission and excitability in the network, initiating an epileptogenic focus and triggering a functional spreading phenomenon for the whole neuronal ensemble, with very disrupting consequences to the brain.

Taken together, our study suggests that iAβo exerts two effects (pre- and post-synaptic) that seem to be site independent for its expression, but both initially triggered signal transduction pathways (Fig.11a). They could provide a potential mechanism to explain early stages of AD, when iAβo accumulates, increasing neuronal hyperexcitability and becoming an epileptogenic focus at the AD onset. Therefore, the prevention of early effects of iAβo, which are mainly intracellular, can potentially be a new therapeutic target for AD. Early treatment could partially or totally reverse the molecular mechanisms by which the peptide initially triggers the synaptic impairment observed in patients in more advanced stages of this disease.

## 5. Conclusions

Taking together *in vitro, ex vivo* and *in vivo* data, we conclude that human and synthetic iAβo induce hyperexcitability through a synaptic-driven mechanism involving AMPA receptors and NO as a retrograde signaling molecule. Additionally, we describe a novel phenomenon in which the functional effect of iAβo on NO production extends to surrounding neurons in a coordinated way. These important findings give new a new meaning to intracellular Aβ accumulation, an early event in the pathology, and uncovers a novel mechanistic explanation for the functional spreading of iAβo-mediated hyperexcitability in a neuronal ensemble.

## 6. Abbreviations

AMPA: α-amino-3-hydroxy-5-methyl-4-isoxazolepropionic acid
APs: Action potentials
APP: Amyloid precursor protein
Aβ: Amyloid-β peptide
CLR: Chelerythrine
EA: Alzheimer’s disease
eEPSCs: Evoked excitatory post-synaptic currents
elPSCs: Evoked inhibitory post-synaptic currents
eNOS: Endothelial nitric oxide synthase
h-iAβo: human AD brain-derived Aβ oligomers
iAβo: synthetic intracellular Aβ oligomers
iNOS: Inducible nitric oxide synthase
mEPSCs: Miniature excitatory post-synaptic currents
mIPSCs: Miniature inhibitory post-synaptic currents
mPSCs: Miniature post-synaptic currents
Nav: Voltage-dependent sodium channel
nNOS: Neuronal nitric oxide synthase
NO: Nitric oxide
NOS: Nitric oxide synthase
PKC: Protein kinase C
PLC: Phospholipase C
PSEN1: Presenilin-1
TTX: Tetrodotoxin
Vm: Membrane potential

## 7. Declarations

### Ethics approval

All experimental procedures were according to Institutional Animal Care and Use Committee at the University of Concepción animal research regulations.

### Consent for publication

Not applicable

### Availability of data and materials

The datasets used and/or analyzed during the current study are available from the corresponding author on reasonable request.

### Author Contributions

EJFP and LGA designed experiments, discussed the results, contributed to all stages of manuscript preparation and editing. Material preparation, data collection and analysis were performed by EJFP, BM, DAB, CP, NORL, MPE, JPM, CF, RB, US. All authors read and approved the final manuscript.

## Acknowledgments

The authors would like to thank Marco Fuenzalida, Juan Pablo Henriquez and Marcela Torrejón for their invaluable comments and for helpful discussion. The authors also thanks Agenor Limon and Lauren Aguayo for having read and provided useful suggestions to the manuscript. For technical support we thank: Laurie Aguayo, César Lara, Daniela Nova, Alejandra Ramírez, Ixia Cid, Javiera Gavilan and Jocelin González.

