## Additional file 1 for "Functional spreading of hyperexcitability induced by human and synthetic intracellular Aβ oligomers"

### 1. Supplementary Methods

1.1. **Preparation of Aβ oligomers.** Briefly, Aβ was dissolved in 1,1,1,3,3,3-Hexafluoro-2-propanol (HFIP) (10 mg/mL) (Merck Millipore, USA) and incubated in a parafilm sealed tube at 37°C for 2 hours. Then, the solution was incubated at 4°C for 20 min and aliquots of 5 µL were placed in 1.5 mL open lid Eppendorf tubes to allow evaporation. Aliquots were stored at -20°C. To obtain an oligomer-rich solution, nanopure water was added to obtain a final concentration of 80 µM and the tubes were incubated at room temperature for 20 minutes. Subsequently, a Teflon-coated magnetic stir bar was added to the solution (size: 2x5 mm) and stirred at room temperature (typically 21ºC) at 500 rpm for 24 hrs. This solution was used to perform the experiments. To characterize the presence of oligomeric Aβ in the preparations used in all the experiments, we used transmission electron microscopy coupled to immunogold staining that showed the presence of spherical or disc-shaped structures of Aβ ranging in sizes from 5-25 nm approximately (supplementary Fig. S2C).

1.2. **Western Blot.** Human brain-derived Aβo was characterized by Western blot analysis. Two different concentrations of Aβo (1 and 0.5 µg of protein) were loaded onto precast NuPAGE 4-12% Bis-Tris gel (Invitrogen) for SDS-PAGE analysis. Gel was subsequently transferred onto nitrocellulose membranes and blocked with 10% nonfat dry milk at 4°C overnight. The membrane was then probed with primary antibodies, A11 (1:1000) and 6E10 (1:6000, BioLegend, USA) diluted in 5% nonfat dry milk for 1 h at RT. HRP-conjugated anti-rabbit IgG and anti-mouse IgG (1:6000, GE Healthcare, USA) were used to detect A11 and 6E10 immunoreactivity, respectively. ECL plus (GE Healthcare, USA) was used to visualize the bands.

1.3. **Atomic force microscopy.** Human brain-derived Aβo was also analyzed by AFM using a non-contact tapping method with a Multimode 8 AFM machine (Bruker, USA). Briefly, 3-4 µl of Aβo was applied onto a fresh-cleaved mica surface and allowed to adsorb at RT overnight. Mica was then washed with 200 µl of deionized water, air-dried and imaged.

1.4. **Immunogold and negative contrast transmission electron microscopy.** Five microliters of Aβo, at a concentration of 50 μM, were applied to carbon-coated Formvar grids (Agar Scientific, UK). Nonspecific immunoreactivity was blocked with 3% bovine serum albumin (BSA) for 30 minutes at room temperature and incubated with the primary antibody anti-Aβ 6E10 (1:50; Novus Biologicals, USA) for 1 hour. A secondary 5-nm gold-conjugated anti-mouse IgG antibody (Merck, Germany) was used at a 1:20 dilution for 30 minutes. Samples were fixed with a 2% glutaraldehyde solution for 5 minutes. Aβo were stained with 5 μL of 0.2 % (wt/vol) phosphotungstic acid and the grid was air-dried. Samples were examined using a JEOL 1200 EX II electronic microscope.

1.5. **Voltage-clamp experiments *in vitro* and *ex vivo*.** To isolate the AMPAergic miniature currents (mEPSCs) *in vivo* and *ex vivo*, synaptic transmission inhibitors were applied using a perfusion system (in μM): 20 DAPV (2-amino-5- phosphonopentanoate), 1 strychnine and 10 bicuculline. The same approach was used to isolate GABAergic miniature currents (mEPSCs) *in vivo* and *ex vivo*, perfusing (in μM): 20 DAPV, 1 strychnine and 20 CNQX (6-Cyano-7-nitroquinoxaline-2,3-dione). All synaptic transmission inhibitors were purchased from Tocris, USA. For recordings of synaptic currents in CA1 hippocampal brain slices (*ex vivo*), the rats were sedated with isoflurane and decapitated. The brain was removed and coronal hippocampal cuts of 300-400 μm thick were made in a VT1200S vibratom (Leica, Germany) in a cold solution containing (in mM): 194 Sucrose, 30 NaCl, 4.5 KCl, 1 $MgCl_2$, 26 $NaHCO_3$, 1.2 $NaH_2PO_4$ and 10 Glucose. Once the slices were obtained, they were allowed to stand in a chamber at room temperature (22 ° C) for 1 hour in artificial cerebrospinal fluid (aCSF) bubbling with 95% $O_2$ and 5% $CO_2$. The aCSF solution contained (in mM): 120 NaCl, 3 KCl, 2 $MgSO_4$, 2.5 $CaCl_2$, 1 $NaH_2PO_4$, 25 $NaHCO_3$ and 20 glucose. The slices were then transferred to the recording chamber with aCSF solution saturated with 95% $O_2$ and 5% $CO_2$ and continuously perfused with oxygenated aCSF at a rate of ~2 ml/min at room temperature (RT). Whole-cell voltage and current clamp recordings were made using an Axopatch 200B amplifier (Axon Instruments, USA) and Digidata 1322A (Molecular Devices, USA). All recordings were filtered at 2.2 kHz and digitized at 10 kHz. Data were acquired using Clampex 10 software (Molecular Devices, USA). Series resistance was continuously monitored and only cells with a stable access resistance were included for data analysis.

1.6. **Current-clamp recordings *in vivo.*** On the day of the recordings, the animals were anesthetized (induction: 3% isoflurane; maintenance: Xylazine/Ketamine 10/100 mg/Kg, supplemented with ketamine 20 mg/Kg). The level of anesthesia was assessed by pinching the foot and by measuring body temperature and respiratory rate. Body temperature was maintained at 37 °C with a thermal blanket (FHC). The animals were fixed in a stereotactic apparatus (SR-6, Narishige, Japan). A local analgesic (lidocaine) was applied as a gel on the stereotaxic system bars to reduce pain during fixation of the head with the stereotaxic system bars, and it was also injected as a liquid under the skin before the first incision. An ophthalmic gel was applied to the eyes to prevent them from drying out during surgery, and the eyes were covered with a piece of cardboard to protect them from light during surgery. The skull was exposed and two small craniotomies (2 mm in diameter) were perforated on both hippocampus (−3.5 mm posterior to bregma; 2.5 mm lateral to bregma) to record in the CA1 area (3 mm deep from the surface of the brain). The Vm of CA1 neurons was recorded in the current clamp mode, using standard techniques for "blind patch" blind cell clamp *in vivo* (99). Before starting the recording of evoked action potentials, a small holding current was applied to stabilize the resting membrane potential (RMP) to -70 mV. The borosilicate electrodes that were used had a resistance of 5-7 MΩ. The internal solution contained (in mM): 135 K-Gluconate, 5.4 KCl, 10 HEPES, 2 Mg-ATP, 0.4 GTP, 0.2 EGTA and 0.2% of biocytin (pH 7.2, adjusted with KOH). The Vm was amplified by an NPI ELC-03XS amplifier (NPI Electronics, Germany) and digitized with a LIH (HEKA Electronik, Germany), using Patch Master software (HEKA Electronik, Germany). Finally, output signals were digitized 1440A Digidata (Molecular Devices, USA) and recorded with Axoscope software (Molecular Devices, USA). 50 Hz noise was removed using a HumBug noise eliminator (Quest Scientific, Canada). For further analysis, only cells with Vm at rest under −55 mV were considered.

1.7. **Histology.** Animals were injected with a ketamine overdose and transcardially perfused with 1x PBS solution followed by 4% paraformaldehyde fixation. The brains were then left in 4% PFA overnight at 4˚C, before washing and storing at 4˚C in PBS. The next day, 50 μm thick coronal slices were *post hoc* processed with the streptavidin method associated with the Cy3 fluorophore (Jackson ImmunoResearch, USA) to visualize neurons containing biocytin. For this, brain slices were incubated with PBS containing 0.3% Triton X-100 (Sigma, Germany), 2% normal goat serum (Thermo Fisher Scientific, USA) and 1:1000 Cy3™ streptavidin (Jackson ImmunoResearch, USA) for 48 - 72 hrs at 4ºC (continuously agitated, and protected from light). After confirming the location of the recorded cell in the hippocampus, slices were blocked with PBST (PBS and 0.3% Triton X-100) plus 7% normal goat serum for 2 hrs at 4 ºC, continuously agitated, and protected from light. Immunostaining was performed using a rabbit anti-Calbindin D-28k antibody diluted 1:1000 (Swant, Switzerland) in a solution containing: PBS, 0.3% Triton X-100, 2% normal goat serum and 1:1000 Cy3™ streptavidin for 24 hrs at 4 ºC (continuously agitated, and protected from light). Slices were washed with PBST (3 times per 10 min at RT, continuously agitated, and protected from light) and then incubated with a secondary Alexa Fluor® 488 Donkey Anti-Rabbit antibody diluted 1:1000 (Jackson ImmunoResearch, USA) using the same protocol and solution of the primary antibody. After washing with PBST (3 times per 10 min at RT, continuously agitated, and protected from light) and PBS (2 times per 10 min at RT, continuously agitated, and protected from light), samples were mounted with Vectashield mounting medium (Vectorlabs, USA). 8 bit images were obtained using a confocal upright Leica TCS SP5 X microscope (Leica, Germany) with a 40x oil immersion objective (1.3 NA) and under the following conditions: for excitation we used 2 laser lines (488 nm, 555 nm) and emission was collected in the 490-540 nm and 569-610 nm ranges, respectively (example in Fig. 8).

1.8. **Simultaneous recordings of electrophysiology and fluorescence**. Hippocampal neurons were incubated with the NO probe DAQ (1,2-aminoanthraquinone) (Sigma, Germany) at a concentration of 2.5 mg/mL for 20 min at 37°C (100, 101). The neurons were washed 3 times with NES and mounted in a well on an inverted microscope (TE200U, Nikon, USA) equipped with a 16-bit IonXEM CCD camera (Andor, Japan), a 20X/0.4 NA objective (Nikon, Japan) and a voltage-clamp configuration for *in vitro* studies. The fluorescent signal for the DAQ probe was obtained by exciting with a bandpass filter (528-553 nm) and collecting the fluorescence with a bandwidth emission filter (590-650 nm) (Nikon, USA). Image acquisition was performed with a computer-controlled Lambda 10-B shutter (Sutter Instruments, USA) using Imaging Workbench 5.0 software (INDEC BioSystems, USA) and exciting for a period of 900 ms at intervals of 1 s during a continuous period of 20 min. Some experiments involved the use of other molecules: L-NAME (NO synthase inhibitor) (Sigma, Germany), 1400W (iNOS inhibitor) (Tocris, USA), SNAP (NO donor) (Sigma, Germany) and CPTIO (NO sequester molecule) (Cayman Chemical, USA). Fresh stocks of all of these reagents were prepared on the same day that the experiment was performed.

1.9. **Data analysis.** Synaptic currents parameters (frequency and amplitude) were analyzed using Mini analysis software (Synaptosoft, Inc., USA), which identifies the currents based on several criteria such as the amplitude the area under the curve and the decay time of each event. As a routine check, we visually inspect all events detected by the software and reject any that did not exhibit the general expected form for synaptic events. Background noise was measured from sections devoid of synaptic events, which oscillated at ~ 2 pA. This value multiplied by 5 (10 pA) was used as a threshold to detect synaptic currents. For spontaneous synaptic recordings, the area under the current trace was integrated (pA · ms) and expressed as charge transferred (nC) during the whole recording (2 minutes) using Clampfit 10.5 (Molecular Devices, USA). For the I/E balance experiments (Fig.3), the analysis was similar, but the baseline current was not included in the analysis. AP parameters were calculated in the first spike of the response as follows: threshold was numerically estimated from first derivative in a V’ versus V phase space projection. From this value, amplitude was calculated to the maximum value reach by the AP. Finally, we obtained the half width of the AP peak expressed as duration. Input resistance was obtained from the slopes in V/I curves in hyperpolarizing current steps. Rheobase was extrapolated from spikes vs. injected current curves using Origin 2019b (Origin Lab, USA). Spontaneous spike firing frequency was obtained using pClamp10 software (Molecular Devices, USA).

### 2. Supplementary Figures

#### Supplementary Figure 1

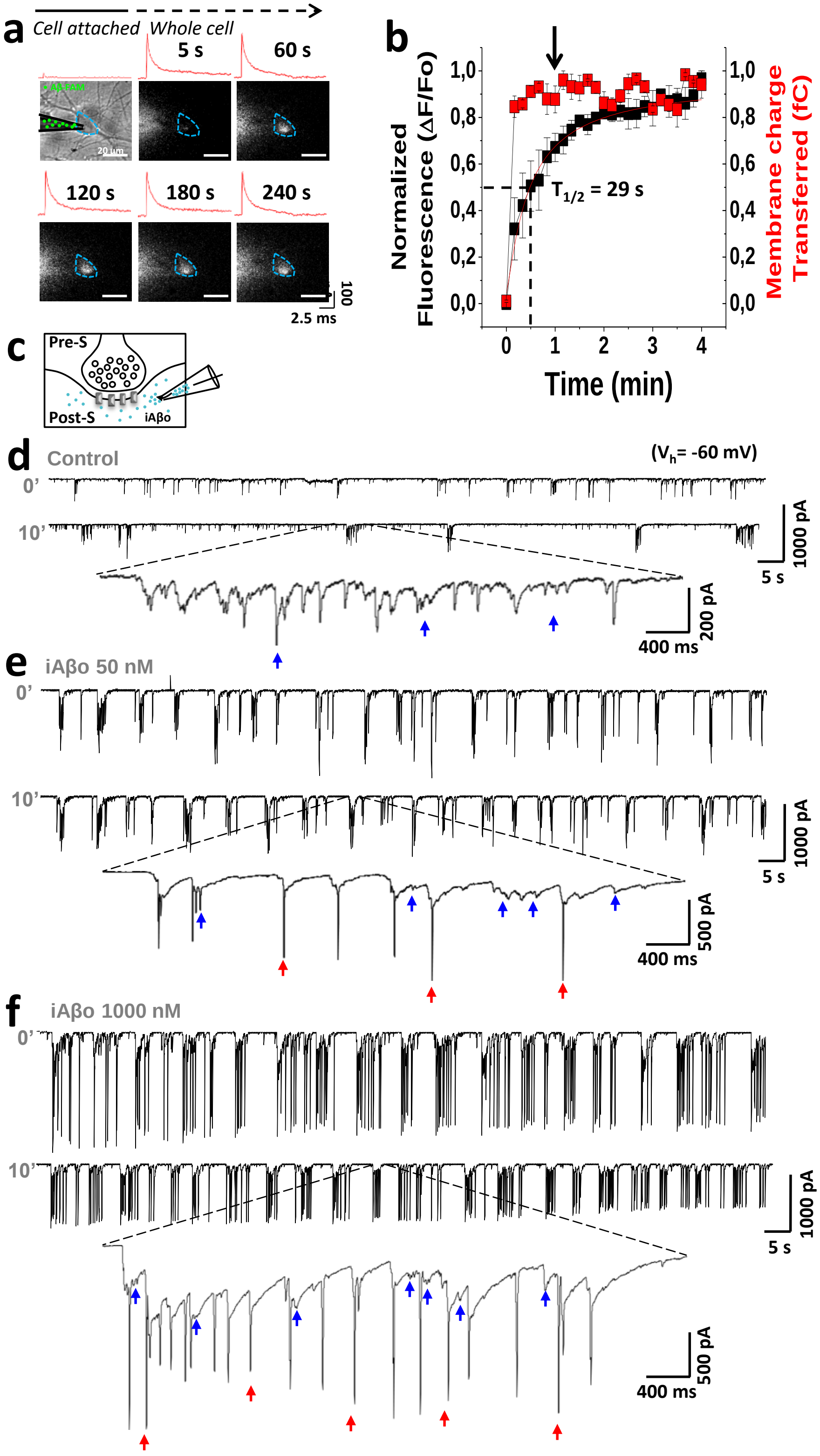

**Supplementary Figure 1. The whole cell technique allows rapid entry of fluorescent Aβo into the intraneuronal compartment. a,** Simultaneous registration of patch clamp and fluorescence showing the entry of fluorescently labeled Aβo (in green) from inside the recording electrode to the intracellular medium. The region of interest (ROI) in (in light blue) delimits the contour of the recorded neuron. Fluorescence quantification was carried out within this region and it was observed that it increases inside the cell throughout the experiment (0-4 min). Along with this, the traces of the capacitive currents recorded at different intervals (in red) are observed. **b,** Quantification of the fluorescence in the ROI previously described, together with the membrane charge transferred. The latter reflects that the solution contained in the patch pipette instantly reaches the intracellular compartment, while the fluorescence accounts for a gradual entry of the peptide, reaching 50% of the total fluorescence value at 29 s ($T_{1/2}$). The black arrow indicates the time at which synaptic currents began to be recorded. **c,** Schematic representation of the synaptic recording, showing the pre-synaptic (Pre-S) and post-synaptic (Post-S) compartment, and the application of iAβo in the latter using the patch electrode (orange squares represent post-synaptic receptors). **d, e, f,** Total synaptic recordings obtained at the beginning (time = 0') and end of the experiment (time = 10'), demonstrating a rapid and marked increase in the frequency and amplitude of synaptic currents as the concentration of intracellular iAβo oligomers (iAβo) increases from 50 nM (**e**) to 1000 nM (**f**) ($V_h$ = -60 mV). Bursts of synaptic currents (arrows in blue) and spikes in current-recording mode are observed in red, which increase as the concentration of iAβo in the recording electrode augmented. Line charts represent the average ± SEM. n = 18 cells per condition.

**Supplementary Figure 2**

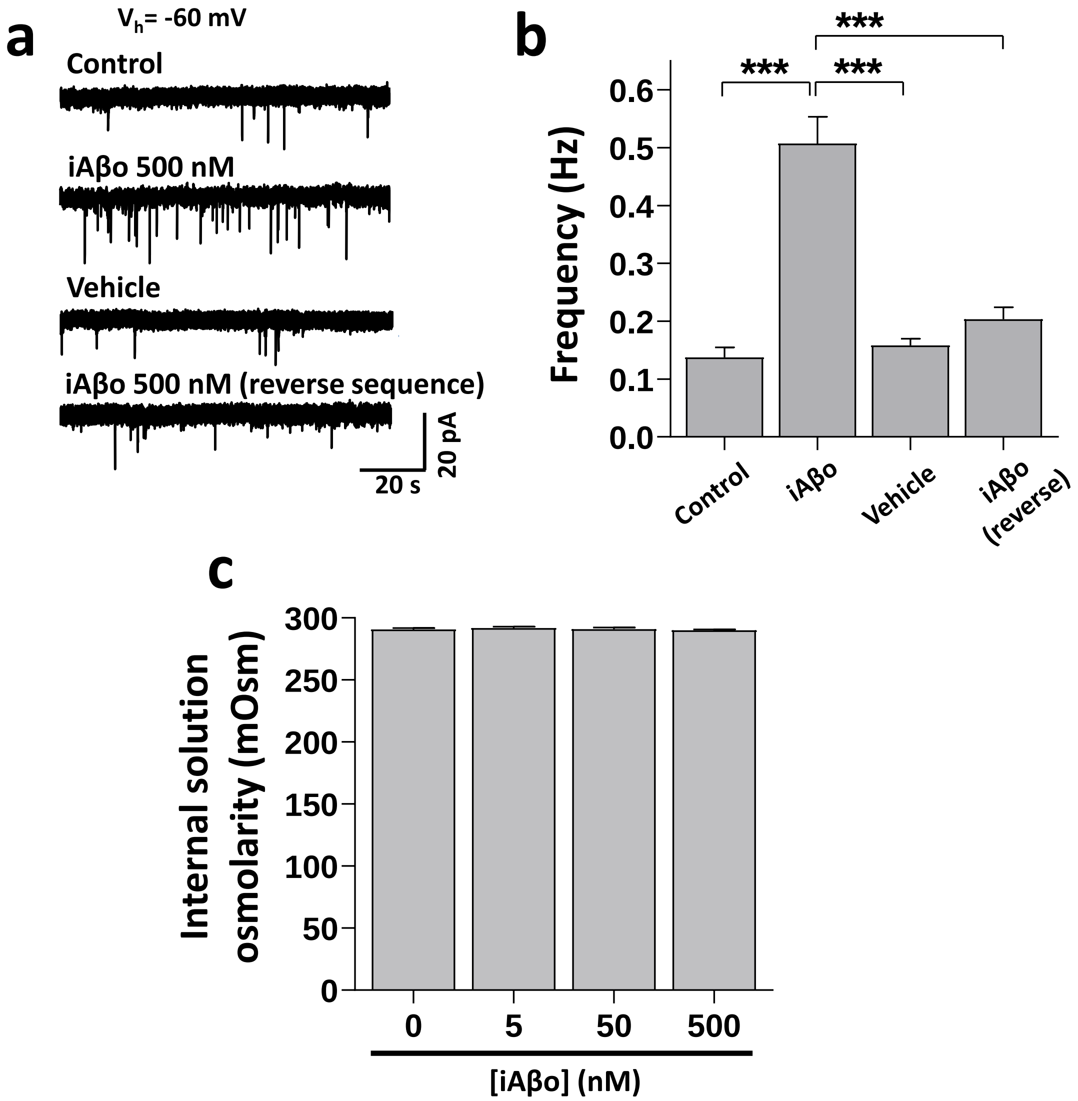


**Supplementary Figure 2. Reverse iAβo do not have an effect on frequency of miniature post-synaptic currents *in vitro*. a, b,** Representative mPSCs traces (**a**) and frequency quantification (**b**) of iAβo, reverse sequence and vehicle controls (One-Way Welch's ANOVA with Games-Howell post-hoc test for: F(3,16.79)=18.78, p=1.30E-5. p-values for post-hoc test: control vs. iAβo 500 nM: 1.07E-4, iAβo vs. vehicle: 2.49E-4 and iAβo vs. iAβo reverse: 4.97E-4). **c,** Internal solution osmolarity measurements show no change when adding different concentrations of iAβo. Bar charts represent the average ± SEM for control (n=9), iAβo (n=9), vehicle (n=9) and iAβo reverse (n=9) cells. *** denotes p <0.001.

**Supplementary Figure 3**

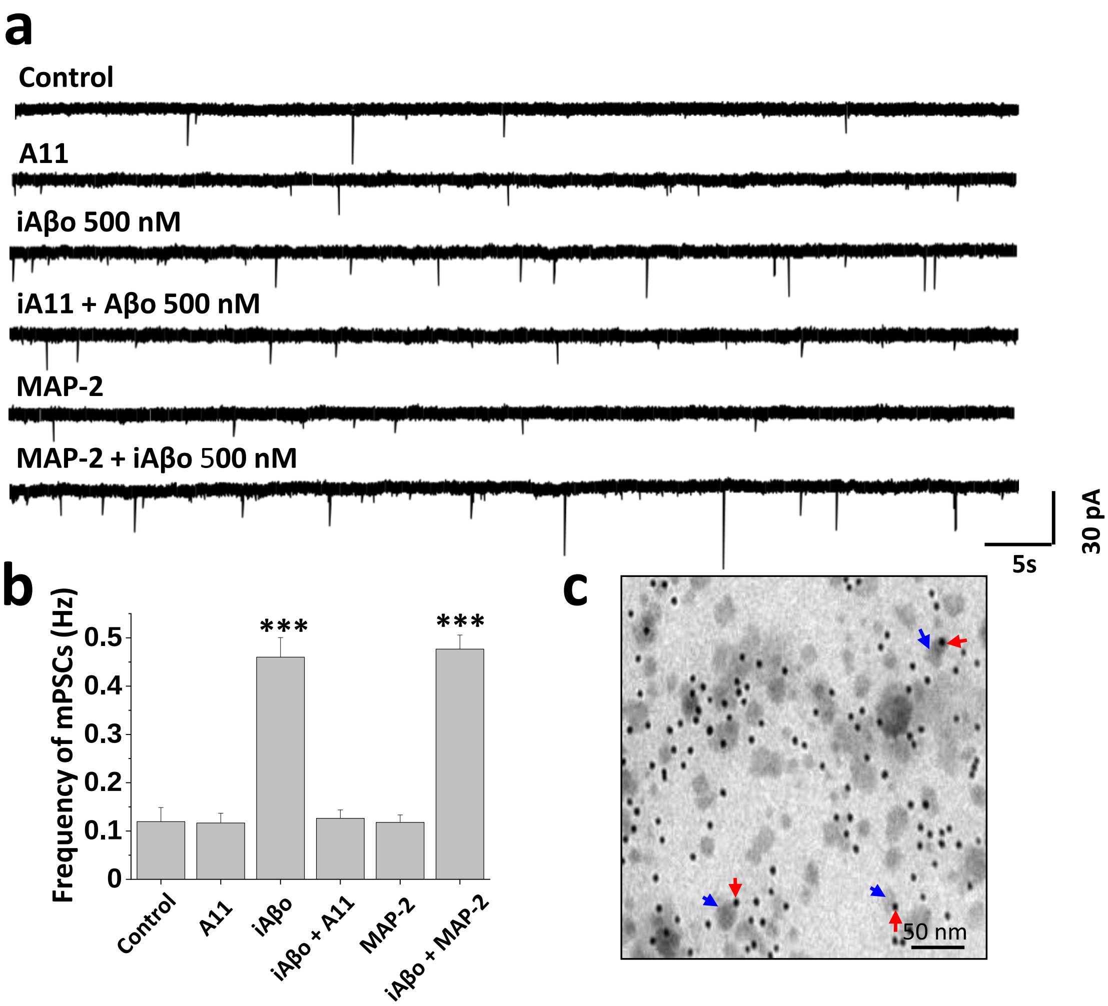


**Supplementary Figure 3. Pre-incubation with A11 antibody attenuates the intracellular synaptic effect of iAβo on the frequency of miniature post-synaptic currents *in vitro*. a,** Representative traces showing the increase in the frequency of mPSCS after application of iAβo 500nM alone or pre-incubated for 10 min with antibody A11 or MAP-2. Intracellular dialysis of the antibodies did not have an effect per se on the frequency of mPSCs ($V_h$ = -60 mV). **b,** Quantification of the frequency of mPSCs under the conditions described in **a**, showing that iAβo increases the frequency of miniature synaptic currents, but this effect is diminished to control levels in a similar way with A-11 pre-incubation. No differences in the effect of iAβo are observed when pre-incubating with an antibody for MAP-2 (one-way ANOVA with Tukey post-hoc: $F(5,44)=27.436$, $p=1.7E-12$).

**c,** Electronic micrographs demonstrating the presence of amyloid oligomeric aggregates (arrows in blue) in the preparations used. Along with that, the presence of gold nanoparticles coupled to the secondary antibody used for Aβ immunodetection is also observed (arrows in red). Bar charts represent the average ± SEM for control (n=8), A11 (n=8), iAβo (n=9), iAβo+A11 (n=9), MAP-2 (n=9) and iAβo+MAP-2 (n=7) cells. *** denotes $p < 0.001$.

**Supplementary Figure 4**

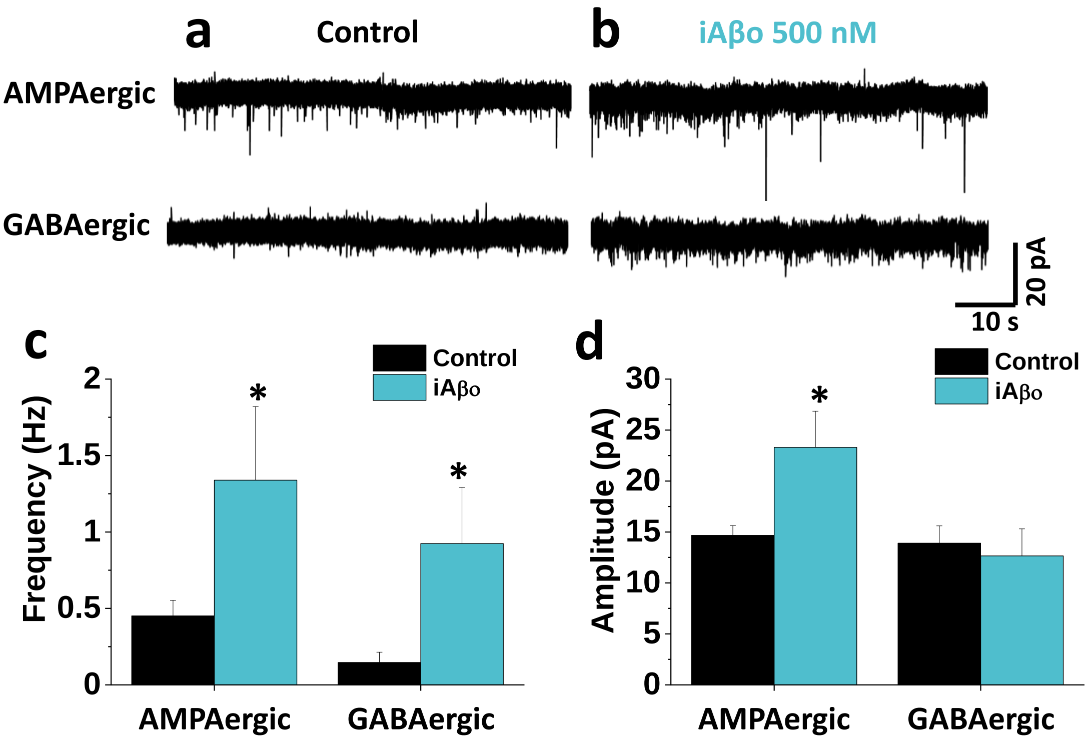


**Supplementary Figure 4. iAβo increases the frequency of AMPAergic and GABAergic synaptic currents in CA1 hippocampal neurons *ex vivo*. a, b,** Representative AMPA and GABA mPSCs obtained in acute hippocampal slices for control condition and with iAβo 500 nM, respectively ($V_h$ = -60 mV). **c,** Quantification of the frequency for AMPAergic (unpaired Student's t-test with Welch's correction: t(6.54)=-2.306, p =3.39E-2) and GABAergic (unpaired Student's t-test with Welch's correction: t(6.40)=-2590, p=1.97E-2) mPSCs. **d,** Amplitude quantification for AMPAergic (unpaired Student's t-test with Welch's correction: t(10.32)=-2.540, p=1.95E-2) and GABAergic (unpaired Student's t-test with Welch's correction: t(10.94)=0.394, p=7.01E-1) miniature currents. Bar charts represent the average ± SEM for control (n=12) and iAβo (n=7) cells. * denotes p <0.05.

**Supplementary Figure 5**

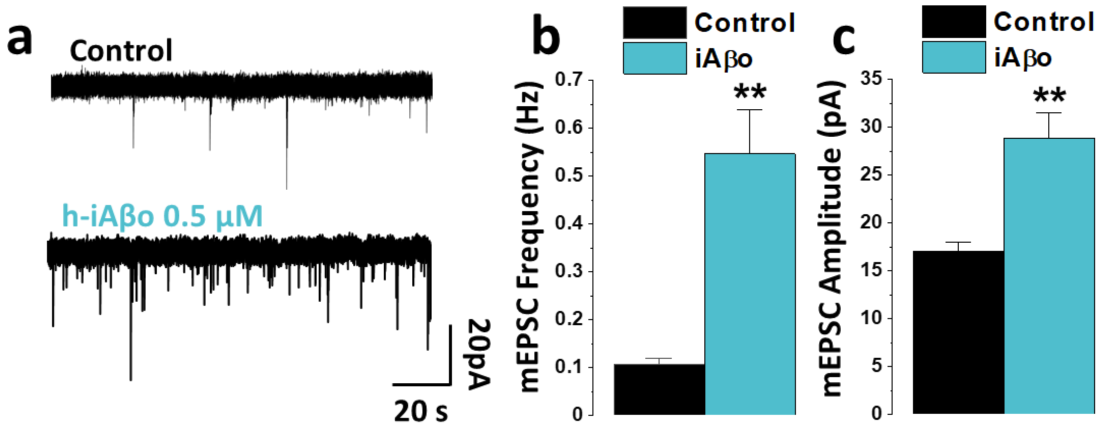


**Supplementary Figure 5. h-iAβo increased AMPA-R mediated mEPSCs *in vitro*. a,** Representative traces of AMPA mEPSCs in control condition and with intracellular application of h-iAβo 0.5 µM ($V_h$ = -60 mV). **b, c,** Quantification of the frequency (**b**) (n=7) (t(7.29)=-4.686, p=2.01E-3) and amplitude (**c**) (n=8) (t(8.73)=-4.323, p=2.03E-3) of the AMPAergic miniature currents, demonstrating a significant increase in presence of h-iAβo. Scatter plots represent the average ± SEM. Unpaired Student's t-test with Welch's correction for (**b**) and (**c**). ** denotes p < 0.005, *** p <0.001.

**Supplementary Figure 6**

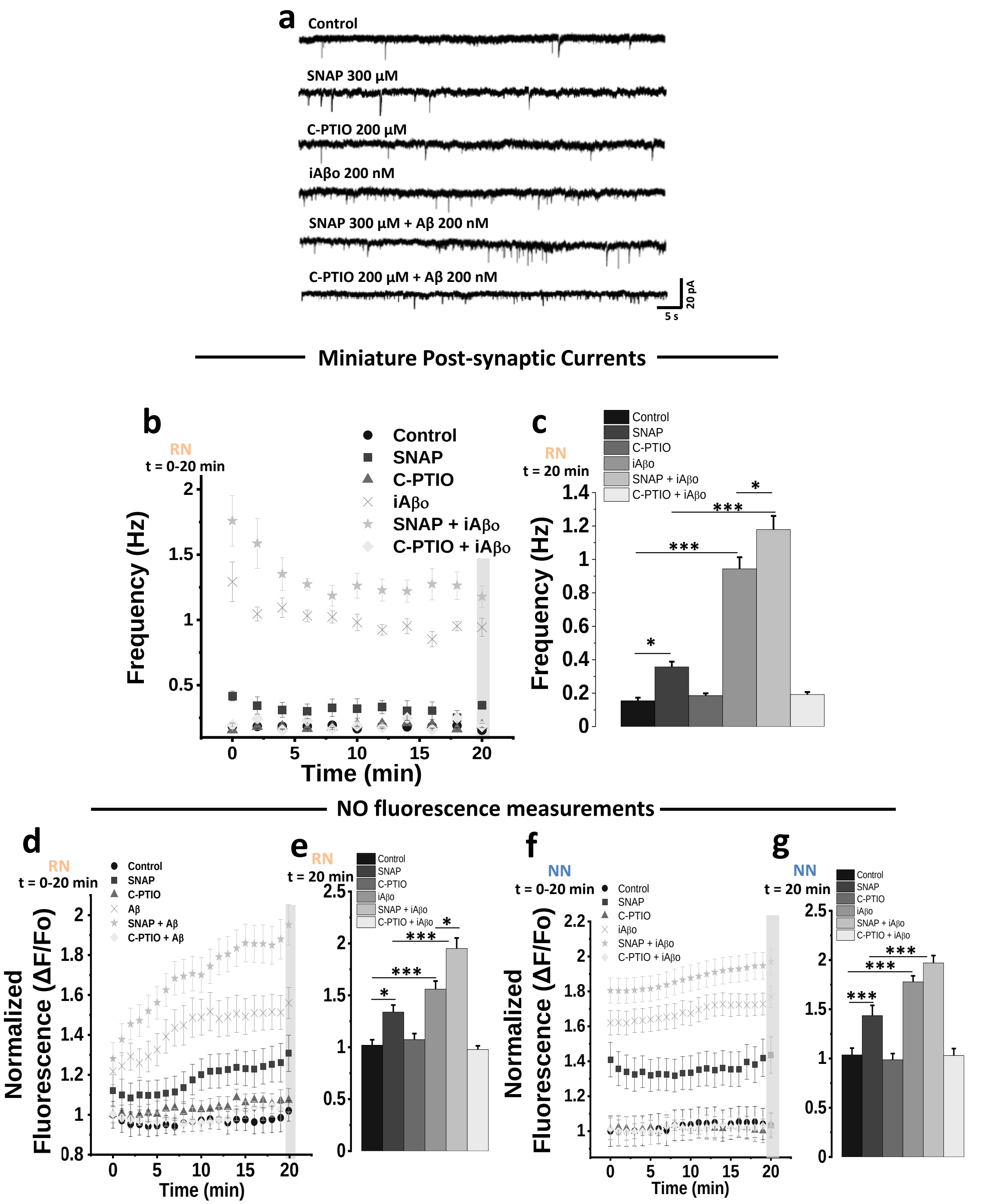

**Supplementary Figure 6. Nitric oxide is involved in the pre-synaptic retrograde signaling of iAβo on the frequency of miniature synaptic currents. a, b, c,** Representative mPSCs traces obtained in absence and presence of Aβ or 200 nM using a NO donor molecule (300 µM SNAP, in gray) or NO scavenger (C-PTIO 200 µM) ($V_h$ = -60 mV). It is observed that SNAP per se has an effect on the frequency of synaptic currents, but when co-applying iAβo+SNAP this effect increases considerably, even exceeding the effect that iAβo has on its own. On the contrary, the application of C-PTIO did not affect the frequency of mPSCs, but co-applied with iAβo decreased the frequency to control levels. The bar graph in (**c**) represent the data recorded at time 20' obtained from graph **b**. **d - g,** Relative levels of NO (expressed as fluorescence) obtained throughout the course of the experiment and at 20' for RN (**d** and **e**) and NN (**f** and **g**). Bar and line charts represent the average ± SEM. Control (n=6), SNAP (n=6), C-PTIO (n=6), iAβo (n=6), iAβo+SNAP (n=6), iAβo+C-PTIO (n=6) for RN and Control (n=57), SNAP (n=51), C-PTIO (n=49), iAβo (n=56), iAβo+SNAP (n=52), iAβo+C-PTIO (n=53) for NN. One-way ANOVA with Games-Howell comparison for (**c**): F(5,30)=89.902, p=3.49E-23. p-values for post hoc test: Control vs. SNAP: 4.09E-02, Control vs. iAβo: 3.17E-13, SNAP vs. SNAP + iAβo: 7.56E-14 and iAβo vs. iAβo + SNAP: 4.29E-2. One-Way Welch's ANOVA with Games-Howell post-hoc test for (**e**): F(5,30)=34.685, p=1.34E-11. p-values for post hoc test: Control vs. SNAP: 4.18E-02, Control vs. iAβo: 1.38E-4, SNAP vs. SNAP + iAβo: 5.84E-7 and iAβo vs. iAβo + SNAP: 2.27E-4. One-Way Welch's ANOVA with Games-Howell post-hoc test for (**g**): F(5,312)=36.376, p=2.57E-29. p-values for post hoc test: Control vs. SNAP: 4.26E-6, Control vs. iAβo: 9.10E-13, SNAP vs. SNAP + iAβo: 2.98E-4. * denotes p <0.05, *** p <0.001.

**Supplementary Figure 7**

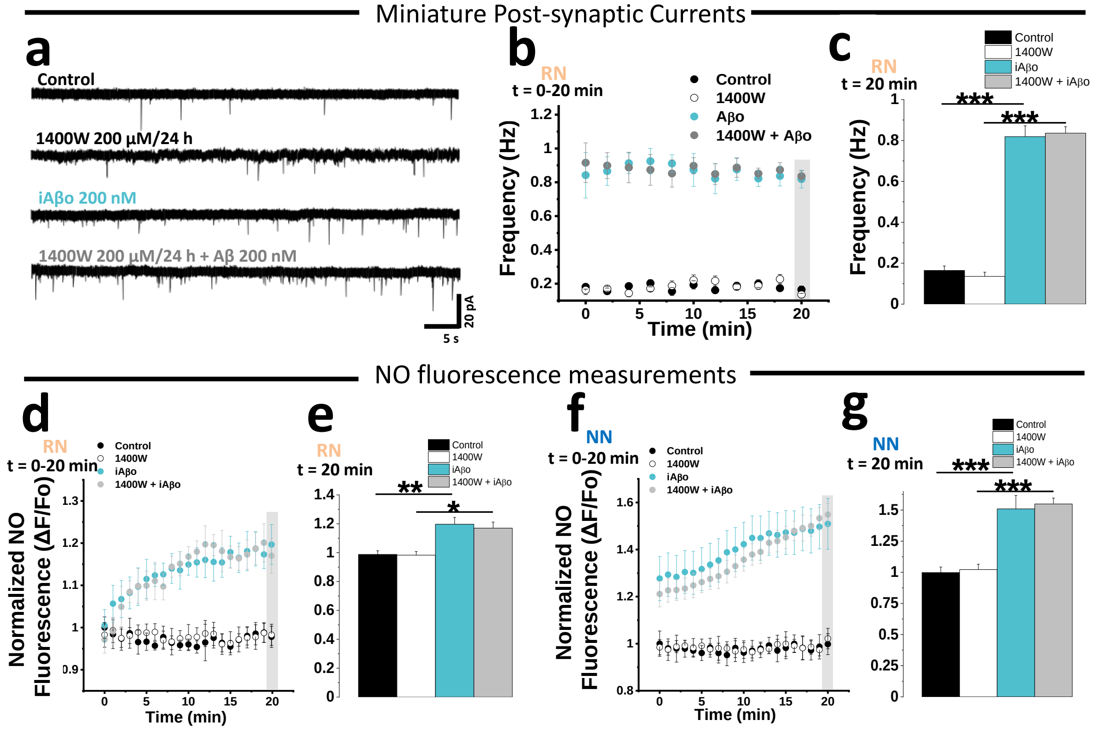


**Supplementary Figure 7. iNOS inhibitor does not affect iAβo actions on synaptic currents frequency. a - c,** Representative recordings and quantification of the frequency of miniature post-synaptic currents in absence and presence of iAβo or 200 nM, together with the co-application of an iNOS inhibitor (1400W) 200 µM for 24 hours ($V_h$ = -60 mV). It is observed that 1400W *per se* does not have an effect on the frequency of synaptic currents. On the other hand, by pre-incubating the culture with 1400W and applying iAβo, it does not change its effect on the frequency of mPSCs. The bar graph in (**c**) was obtained from the data recorded at time 20 '. **d – g** NO fluorescence recordings obtained from the recorded neuron (RN) (**d** and **e**) and from adjacent neurons (NN) (**f** and **g**). Both, RN and NN cells, exhibit an increase in NO when applying iAβo in the RN neuron. This effect does not change significantly when iAβo is applied in a culture that has been pre-incubated with iNOS inhibitor. Line and bar graphs represent the average ± SEM for control (n=8), 1400W (n=7), iAβo (n=9) and iAβo + 1400W (n=10) for RN and control (n=78), 1400W (n=82), iAβo (n=75) and iAβo + 1400W (n=79) for NN. One-Way Welch's ANOVA with Games-Howell post-hoc test for (**c**): F(3,30)=28.474, p=6.50E-9. p-values for post hoc test: Control vs. iAβo: 9.70E-7,

1400W vs. iAβo + 1400W: 3.52E-6. One-Way Welch's ANOVA with Games-Howell post-hoc test for (**e**): F(3,30)=9.0002, p=2.10E-4. p-values for post hoc test: Control vs. iAβo: 2.41E-3, 1400W vs. iAβo + 1400W: 1.02E-2. One-Way Welch's ANOVA with Games-Howell post-hoc test for (**g**): F(3,310)=21.348, p=1.35E-12. p-values for post hoc test: Control vs. iAβo: 4.99E-7, 1400W vs. iAβo + 1400W: 8.97E-8. * denotes p <0.05, ** p <0.005, *** p <0.001.

**Supplementary Figure 8**

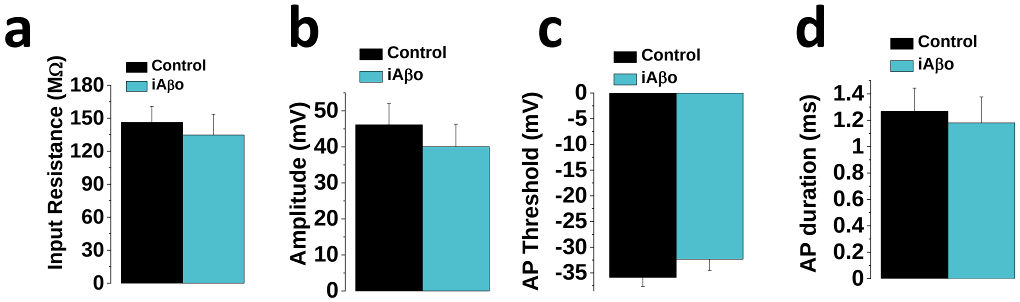


**Supplementary Figure 8. iAβo does not change intrinsic excitability membrane parameters in hippocampal neurons *in vivo*. a – d** Quantification of input resistance and AP parameters: amplitude, duration (half-width) and threshold, all of which do not show significant differences between the conditions tested. Bar and line charts represent the average ± SEM for control (n=10) and h-iAβo (n=6) cells of at least 6 rats.

**Supplementary Figure 9**

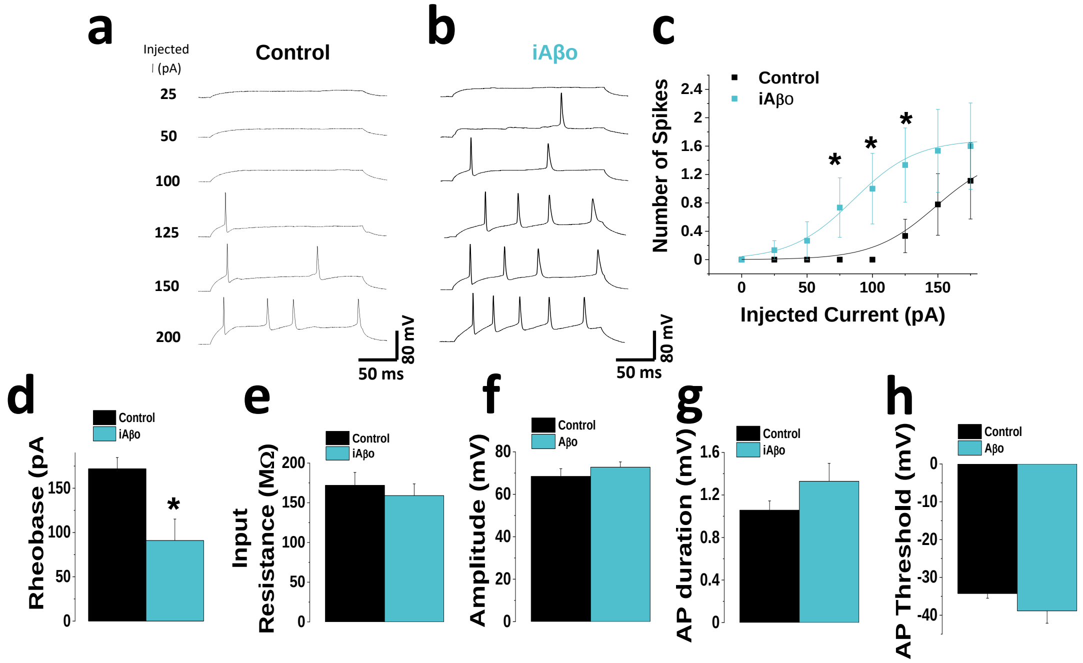


**Supplementary Figure 9. iAβo increased the firing of action potentials evoked by current injection in hippocampal neurons *in vitro*. a, b,** Hippocampal neuron action potential (AP) recordings in the absence (**a**) and presence of 500 nM iAβo (**b**). **c,** Relationship between the number of triggered AP and the injected current intensity for the experimental conditions described previously. **d,** Rheobase constant decreased for iAβo condition. **e – h,** Quantification of input resistance and AP parameters: amplitude, duration (half-width) and threshold, all of which do not show significant differences between the conditions tested. Bar and line charts represent the average ± SEM. n=12 cells per condition. * denotes $p < 0.05$.

**Supplementary Figure 10**

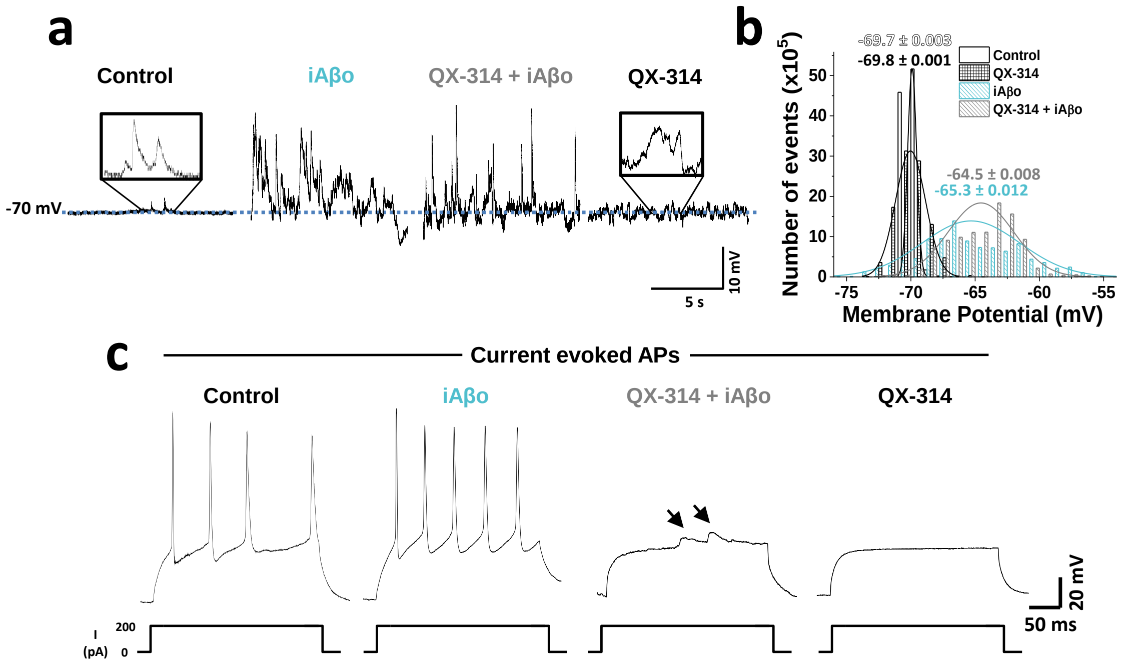


**Supplementary Figure 10. Intracellular blockade of voltage-regulated $Na_v$ channels does not prevent depolarization of the membrane activated by iAβo. a,** Representative recordings obtained without current injection, showing membrane potential (Vm) fluctuations under the different conditions tested. Small variations in the value of Vm are observed for control condition, which are exacerbated in the presence of iAβo 500 nM, while the co-application of iAβo with QX-314 did not diminished the intracellular effects of iAβo on Vm fluctuations. QX-314 by itself did not show any differences with respect to control conditions. **b,** Histogram showing the distribution of Vm values along with average values ± SEM in the different experimental conditions shown in A. **c,** Current injection experiments demonstrating that, under the control and iAβo conditions, the generation of action potentials was not inhibited, while $Na_v$ intracellular block by QX-314 prevented spiking of neurons with and without iAβo. Black arrows indicate that even when $Na_v$ was effectively blocked, depolarizing post-synaptic potentials were appreciated when iAβo was present. This did not occurred for the condition with QX-314 alone. n=10 cells per condition.

**Supplementary Figure 11**

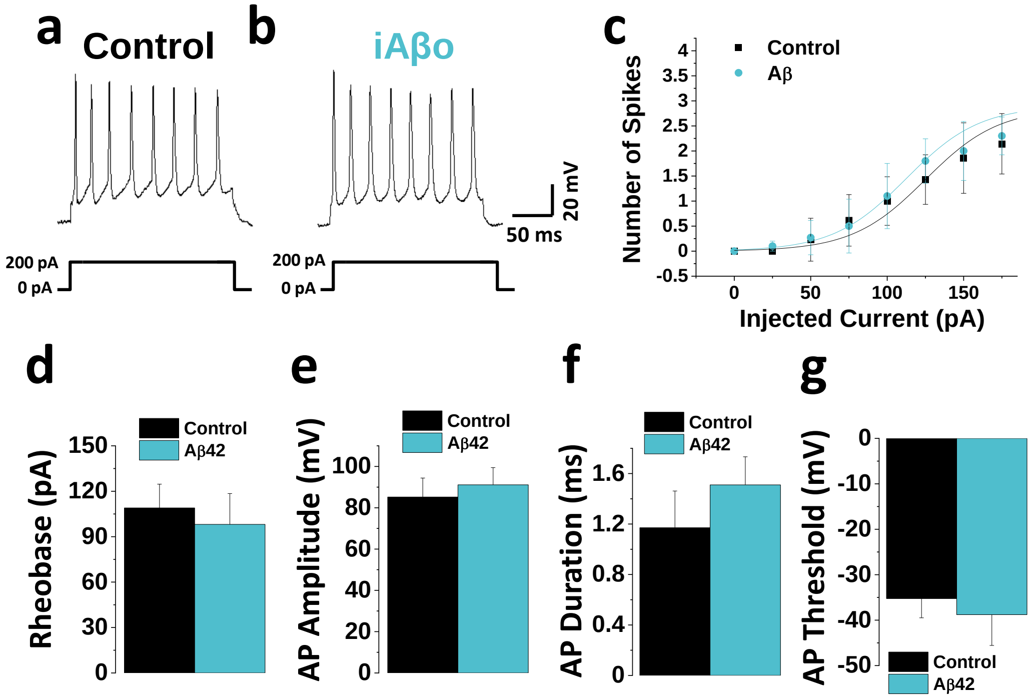


**Supplementary Figure 11. iAβo did not increase the firing of action potentials in dorsal root ganglion neurons (DRG) *in vitro*. a, b,** Representative current-evoked action potentials in the absence (**a**) and presence of 500 nM iAβo (**b**). **c,** Relationship between the number of evoked action potentials and the injected current intensity for the experimental conditions described in **a** and **b**. **d – g,** Quantification of the rheobase constant (**d**) and AP parameters amplitude (**e**), duration (**f**) (expressed as half-width) and threshold (**g**) showing no statically differences between both conditions. Bar and line charts represent the average ± SEM. n=10 cells per condition.
